# The cellular diversity of human cerebrospinal fluid following intraventricular hemorrhage revealed by single-nucleus RNA sequencing

**DOI:** 10.1101/2025.06.05.657834

**Authors:** Sonia Malaiya, Riccardo Serra, Marcia Cortes-Gutierrez, Bradley E. Wilhelmy, Emily Jusuf, Mayur Somalinga, Dave Peprah, Hirschel Nambiar, Kevin T. Kim, Jordan R. Saadon, Parantap D. Patel, Steven K. Yarmoska, Maureen Rakovec, John Kim, Catherine Lei, Shreyas Panchagnula, Reina Ambrocio, Meagan Cherney, Nicholas A. Morris, Natarajan Ayithan, Xiaoxuan Fan, Volodymyr Gerzanich, J. Marc Simard, Neeraj Badjatia, Gary Schwartzbauer, Gunjan Y. Parikh, Seth Ament, Prajwal Ciryam

## Abstract

**Background:** Intraventricular hemorrhage (IVH) is a common and severe complication of hemorrhagic brain injury. Current treatments offer limited improvement in long-term neurological outcomes. Inflammatory responses in the cerebrospinal fluid (CSF) after IVH are thought to drive secondary injury, but the cellular mechanisms underlying this inflammation remain poorly defined.

**Methods:** We performed single-nucleus RNA sequencing of leukocytes isolated from CSF collected through external ventricular drains in subjects with intracerebral (*n* = 6) or subarachnoid (*n* = 1) hemorrhage. We characterized transcriptionally distinct subpopulations of neutrophils, monocytes, and lymphocytes by comparison to reference datasets. Cell–cell signaling networks were analyzed to infer cytokine-mediated communication, and a flow cytometry panel was developed to validate transcriptomic findings in independent CSF samples.

**Results:** We obtained 11,191 high-quality nuclei comprising neutrophils (53.8%), monocytes (26.1%), lymphocytes (17.8%), and non-immune cells (2.4%). Neutrophils segregated into Nascent, Quiescent, and Interferon-Activated states. Monocytes exhibited classical phenotypes that include interferon-activated states (characterized by expression of *VCAN* or *PROK2*) and CXC-chemokine expressing states (characterized by expression of *CXCL5* or *CXCL8*). Lymphocytes were mainly naïve and central memory CD4⁺ T cells. Cell–cell signaling analysis predicted strong CXC chemokine signaling from monocytes to neutrophil subsets and IL-1 family–driven inflammatory responses across multiple populations. Type I and III interferon signaling defined a neutrophil population not previously described in the central nervous system.

**Conclusion:** This study delineates the diverse cellular immune landscape of CSF after IVH. Transcriptomic profiles reveal interferon, IL-1, and CXC chemokine signaling networks as potential therapeutic targets to mitigate secondary injury.

## Introduction

Bleeding into the cerebral ventricles, known as intraventricular hemorrhage (IVH), is an ominous complication of hemorrhagic brain diseases, including intracerebral hemorrhage (ICH), subarachnoid hemorrhage (SAH), and traumatic brain injury (TBI). In adults, it results in death or severe disability in over 50% of cases^1^. The development of elevated intracranial pressure and hydrocephalus contribute to morbidity and mortality^2–4^. Treatment options are limited, and while around 50% of patients require temporary drainage of cerebrospinal fluid (CSF) with an external ventricular drain (EVD)^5,6^, lifelong CSF shunting via ventriculoperitoneal shunting is needed in up to 40% of subjects^7,8^. Intraventricular injection of tissue plasminogen activator does not improve functional outcomes or reduce the rate of shunting^7^.

As new prognostic and treatment approaches are sought for this morbid disease, promising strategies may arise from the critical role of inflammation in IVH. The deposition of blood into the cerebral ventricles stimulates a profound inflammatory response that recruits leukocytes into the central nervous system^9,10^. In the first several days after injury, this response is dominated by myeloid cells: first neutrophils, then monocytes and macrophages. These cells interact with CNS resident cells, with both pathological and reparative consequences. On one hand, inflammation after IVH can cause neuronal injury, cerebral edema, and hydrocephalus; on the other, it may stimulate neurogenesis and clearance of hemorrhagic debris^11–14^. Because of the numerous and pleiotropic contributors to inflammation in IVH, comprehensive approaches are needed to characterize this process.

Here, we sought to characterize the CSF inflammatory response in patients after IVH, using single-nucleus RNA sequencing. Our approach identified distinct states of neutrophils and monocytes in the CSF, including a population of interferon-induced neutrophils not previously described in the central nervous system. These findings illustrate the complex cellular inflammatory response to IVH and point to potential novel therapeutic targets for further investigation.

## Results

### A snRNA-seq atlas for cellular diversity of ventricular CSF in the context of intraventricular hemorrhage

Previous scRNA-seq studies of CSF samples from patients with IVH did not capture its full cellular diversity. Based on prior CSF cell count data in IVH^15^, we anticipated that the CSF samples we collected would be highly represented in neutrophils and other fragile populations that could be depleted during sample processing. We reasoned that single-nucleus RNA sequencing (snRNA-seq) could provide a means to limit these processing-induced biases and might yield a more representative cellular atlas of IVH. We obtained ventricular CSF of seven study participants who presented with IVH, either directly to the University of Maryland Medical Center or after emergent transfer from another institution (**Fig. 1A**; **Table 1**). Six subjects presented after ICH, with etiologies including hypertension and coagulopathy/anticoagulation-related hemorrhage, with one subject whose etiology was unknown. One subject presented with aneurysmal SAH. Of the seven subjects, five presented with severe deficits in level of consciousness as quantified by Glasgow Coma Scale (GCS) between 3-8 and two presented with a mild deficit (GCS 13-15). Standardized scales of disease severity were consistent with moderate to severe disease. For subjects with ICH, hemorrhages originated in deep structures of the brain in a majority of cases. All subjects had IVH, either at presentation or subsequently in their hospital course. When CSF is obtained for clinical purposes, excess fluid that would otherwise be discarded is sometimes available. We collected this CSF immediately after it was aspirated. As these are convenience samples, they were obtained at a range of time points after disease onset. Non-contrast CT scans of the head from these subjects demonstrate the presence of IVH and show the EVD tip in the lateral ventricles near the foramen of Monro (**Fig. 1B**). CSF is therefore sampled directly from the ventricles, unlike lumbar puncture, which samples CSF from the lumbar cistern in the lower back. CytoSpin cytology data available from the electronic health record for sample aliquots from five of the seven patients indicated extensive infiltration of leukocytes into the CSF, primarily comprised of neutrophils, monocytes/macrophages, and T cells (**Fig. 1C**).

**Figure 1.**
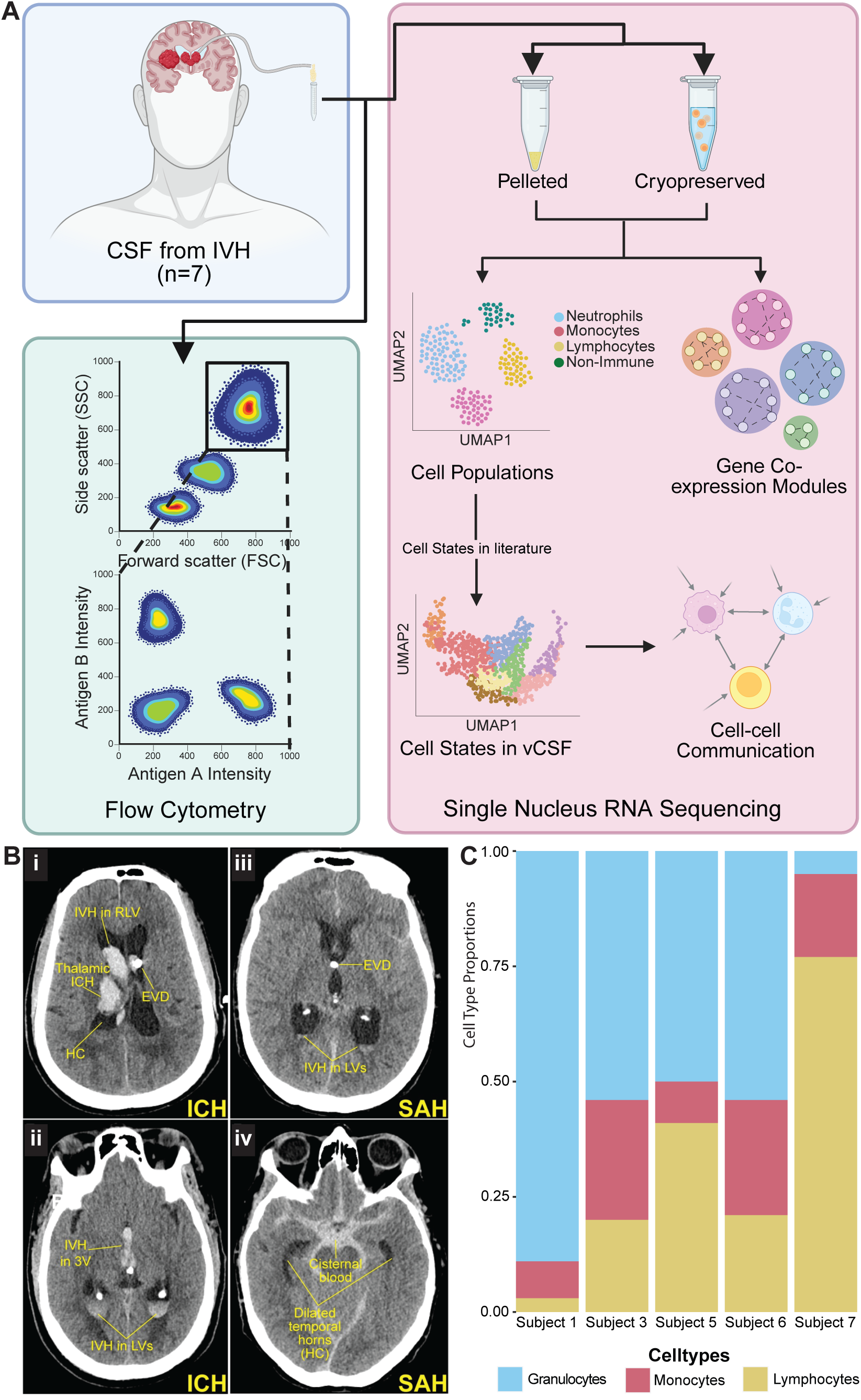
Clinical characteristics of study participants. A. Study design (Created in https://BioRender.com). B. Computed tomography (CT) scans. Axial images (slice thickness=3 mm) from the first CT study obtained after EVD placement for a subject with ICH (i,ii) and SAH (iii, iv). i. Thalamic ICH with IVH most prominent in right lateral ventricle (RLV) and ventriculomegaly consistent with hydrocephalus (HC), with EVD visible in left lateral ventricle. ii. More caudal slice from same CT as in (i) showing IVH in the third ventricle (3V) and bilateral lateral ventricles (LVs). iii. Aneurysmal SAH with small amount of IVH in bilateral LVs, with EVD visible at the foramen of Monro. iv. More caudal slice from same CT as in (iii) showing blood in the basal cisterns in classical aneurysmal SAH pattern, with dilated temporal horns consistent with HC. C. Cell proportions in ventricular CSF assessed by cytospin columns.

snRNA-seq of CSF from these seven study participants yielded 11,191 high-quality nuclear transcriptomes (**Fig. 2A, B**). Neutrophils (n = 6,017; *S100A8^+^*, *CXCR1^+^*, *FCGR3A*^+^, *FCGR3B*^+^) were the most abundant cell type, followed by monocytes (n = 2,920; *VCAN^+^, CXCL8*^+^, *CCL2L1*^+^), and lymphocytes (n = 1,990; *CD247*^+^) (**Table S1**). We also identified a small number of cells expressing glial or neuronal markers (n = 264; e.g., *PTPRD, RSPO3, FLT1, CLDN10, RYR2, RBFOX1, DLG2*) (**Fig 2C**). These cells may derive from damage to the cerebral grey matter during or after the placement of the EVD and were not analyzed further. Cell type proportions identified by snRNA-seq were in the same range as those obtained using CytoSpin columns (**Fig. 1C**), although our pooling strategy does not allow us to match individual samples across the two technologies. Overall, our data represent a relatively unbiased atlas for the cellular diversity of the CSF in the context of IVH.

**Figure 2.**
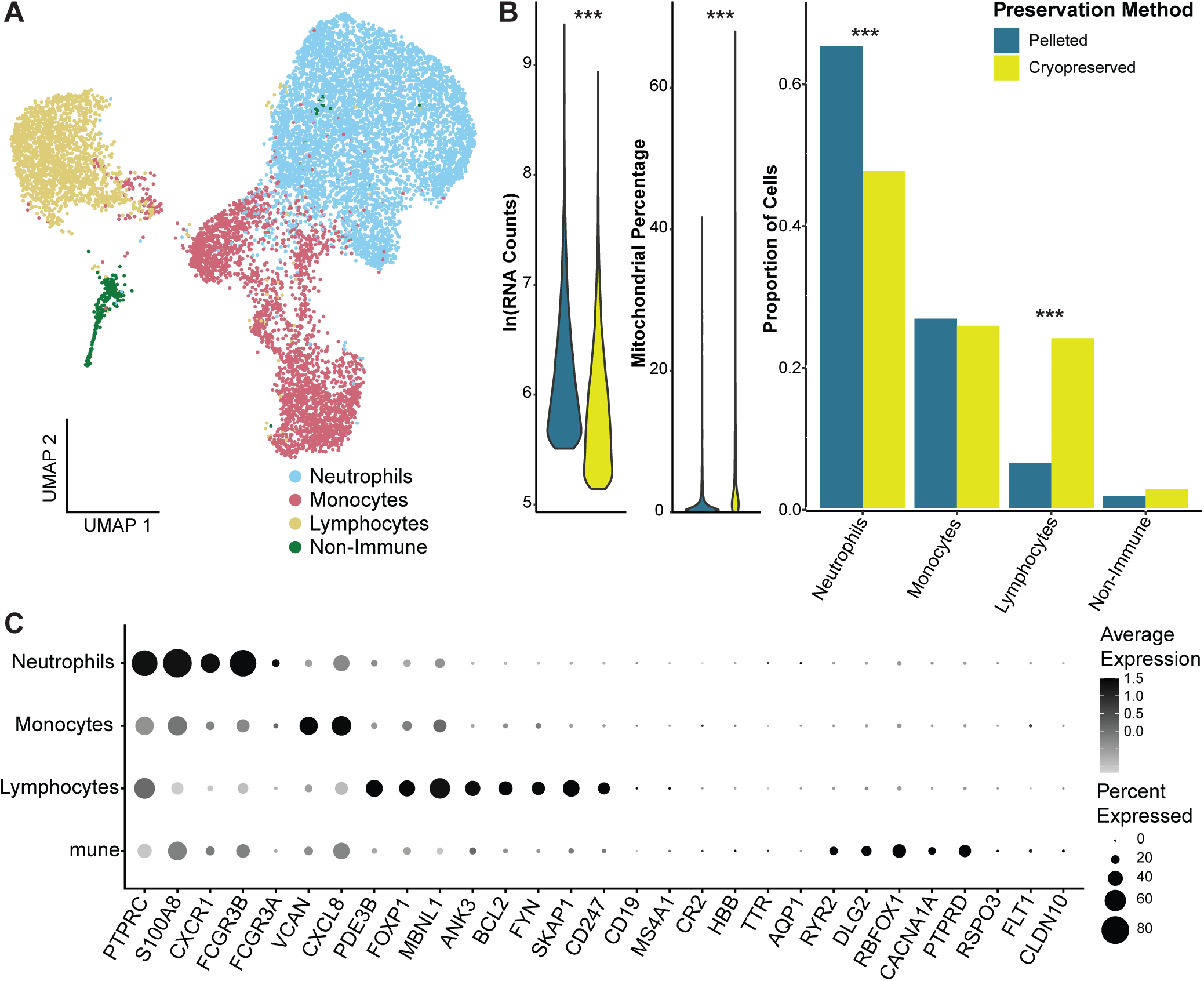
snRNA-seq of ventricular CSF after IVH. A. Uniform manifold approximation and projection (UMAP) plot for snRNA-seq of ventricular CSF (11,191 nuclei from seven study participants). B. Quality metrics comparing cell processing methods. C. Dot plot showing expression of canonical marker genes across cell types.

Participants’ cells were preserved for snRNA-seq processing by two different methods, comparing distinct cell processing protocols in which nuclei were isolated directly from frozen cell pellets vs. cells that had been cryopreserved with DMSO and briefly thawed. snRNA-seq of nuclei derived from pelleted vs. cryopreserved cells yielded a larger number of nuclei from the cryopreservation method, but this method also produced lower RNA counts per nucleus and higher mitochondrial counts per nucleus (**Fig. 2B, Fig. S1 A-C**). The library derived from cryopreserved cells contained a smaller fraction of neutrophils and larger fraction of lymphocytes. It is unclear whether these differences correspond to biological variation among the participants, or whether they reflect a greater vulnerability of neutrophils during cryopreservation and thawing. Nevertheless, these data suggest that all of the cell types in the CSF can be obtained for snRNA-seq by either cell processing strategy.

### Neutrophil diversity in the ventricular CSF

Next, we sought to characterize leukocyte populations in greater detail. Re-integration of the 6,017 nuclei identified as neutrophils revealed eight transcriptionally-distinct neutrophil sub-clusters. All eight of these sub-clusters were detected in both preservation methods (**Fig. S2A-D**), but with varying proportions. We annotated these neutrophil sub-clusters via comparisons to scRNA-seq of neutrophil states in the blood and in lumbar CSF^16^, enabling us to merge the eight sub-clusters into three broad neutrophil populations (**Fig. 3A, Table S4**). The majority of neutrophils form a continuum from Nascent Neutrophils (21.29%) (neu4, neu6, neu8), which expressed *S100A8*, *S100A9*, and *SELL*, to Quiescent Neutrophil states (65.43%) (neu1, neu2, neu5, and neu7), which expressed markers such as *MALAT1* and *CXCL8* (**Table S2**). In addition, a third population expressed genes that are activated during interferon (IFN) responses (13.28%) (neu3), including *IFIT1*, *IFIT2*, *IFIT3*, and *FCGR3B*. These IFN-Activated Neutrophils are a cell population with diverse but not fully characterized functions^17,18^. To our knowledge, they have never been described in the CSF previously.

**Figure 3.**
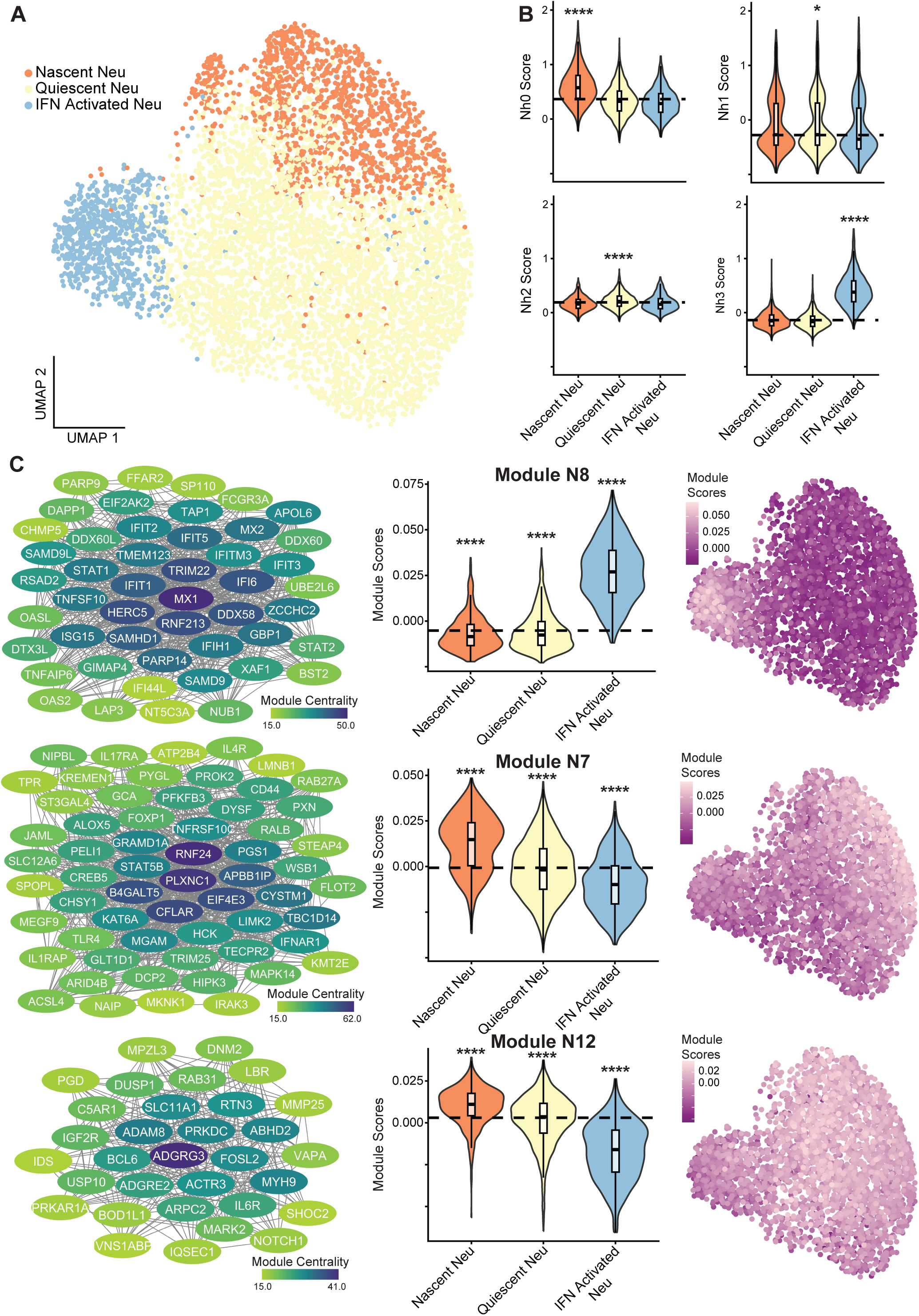
Neutrophil diversity in CSF after IVH. A. UMAP indicating neutrophil subtypes. B. Annotation of neutrophil subtypes by scoring with marker genes from neutrophil subtypes identified in the blood^16^. C. Gene co-expression modules differentially expressed across neutrophil subtypes. Wilcoxon rank-sum test (one-sided for reference marker scoring, two-sided for gene co-expression networks) with Bonferroni correction used for statistical significance testing (see Tables S3, S4). *p < 0.05, **p < 0.01, ***p < 0.001, ****p < 0.0001.

To further characterize these neutrophil populations, we scored them on the activity of marker genes for neutrophil populations identified in the blood in two previous scRNA-seq studies by Wigerblad et al.^16^ (**Fig. 3B, Tables S3, S8**) and Kwok et al.^19^ (**Fig. S3A, Tables S3, S8**). We also scored neutrophils for the activity of 12 gene co-expression modules (N1-N12) computed within the neutrophils from our snRNA-seq data to describe distinct patterns of activity across the neutrophil populations (**Fig. 3C, S4A Table S4**).

Nascent Neutrophils were defined by elevated expression of markers for the “Nh0” immature neutrophil population from the Wigerblad dataset (adjusted p = 2.86×10^-153^), as well as markers for two immature neutrophil populations from the Kwok dataset (S100A8-9 hi neutrophil markers (adjusted p = 6.10×10^-219^) and IL1R2^+^ immature neutrophil markers (adjusted p = 2.55×10^108^). Nascent Neutrophils also displayed high expression of gene co-expression module N7 (adjusted p = 7.27×10^-99^). Genes in this module were over-represented for NF-κΒ activation genes (GO:0051092) and signaling pathways for pattern recognition protein receptors (GO:0002221), Toll-Like receptors (GO:0002224), and cytokine receptors (GO:0019221, GO:0004896), including cytokine-signaling pathways related to IL-12 (GO:0032655), and IL-1 (GO:0032651, GO:0005149).

Quiescent Neutrophils were defined by the expression of long noncoding RNAs that modulate with inflammatory responses (e.g., *MALAT1*, *NEAT1*, and *LUCAT1*^20–23)^ and genes that are associated with neutrophil migration and chemotaxis (e.g., *SSH2*, *DOCK5*, *CXCL8*^24–26)^. While markers of the Wigerblad ‘Nh2’ “transcriptionally inactive” cluster had overall low expression in our dataset, Quiescent Neutrophils had a modestly increased expression of these markers relative to the other neutrophil populations (log_2_FC = 0.190, adjusted p = 4.94×10^-37^). Several gene co-expression modules were upregulated in Quiescent Neutrophils, including N5 (adjusted p = 3.02×10^-16^), N3 (adjusted p = 1.61×10^-39^), N10 (adjusted p = 1.24×10^-27^), and N11 (adjusted p = 1.88×10^-64^). *MALAT1* and *NEAT1* were members of N5, suggesting a role for this module in the epigenetic regulation of inflammatory responses. N5, N3, and N11 were over-represented in genes associated with serine/threonine kinase activity (GO:0004674), with the strongest upregulation of *SIK3*, *STK4*, and *CAMK1D*. Serine/threonine kinases, including *STK4* and *CAMK1D*, have been reported to play a variety of roles in neutrophils^27–29^, but their functional significance in the Quiescent Neutrophil population is unclear. Interestingly, *CAMK1D* and *MALAT1* have also been found to be co-expressed in a subset of granulocytes in the blood^30^.

The neutrophil module N12 consists of several genes that play roles in cell adhesion and migration (ADGRG3, ADGRE2, MYADM, ADAM8). The elevated expression of this module in nascent (adjusted p = 5.03×10^-74^) and quiescent neutrophils (adjusted p = 9.55×10^-06^) may suggest an active recruitment of these cells at the time of sample collection.

IFN-Activated Neutrophils were defined by elevated expression of many canonical interferon-stimulated genes, genes. Markers of the ‘Nh3’ “IFN inducible” neutrophil population in the Wigerblad dataset were strongly induced (adjusted p < 2.23×10^-308^), as was gene co-expression module N8 (adjusted p = 1.31×10^-269^), whose hub genes included canonical interferon-stimulated genes such as *IFI6, IFI30, IFI44*, *IFI44L, IFIT1, IFIT2* and *IFIT3*. Other marker genes annotated as activated in response to symbionts (GO:0140546), viruses (GO:0051607), cytokines (GO:0034097), and interferons (GO:0034341, GO:0034340, GO:0035456) included *MX1, IFITM3, IFIT5, IFI6, CD74, CXCR4, and SELL*. This cell state also had decreased expression of the *IL1B*-containing module N10 (adjusted p = 1.24×10^-63^), consistent with the suppressive effects of interferons on IL-1 production^31^.

### Monocyte/Macrophage diversity in the ventricular CSF

We performed similar analyses to characterize monocyte lineages in ventricular CSF (**Fig. 4**). By sub-clustering the 2,920 monocyte-lineage cells, we identified 6 transcriptionally distinct Seurat clusters annotated based on gene expression markers in the literature as classical monocytes (four clusters, 90.6%), macrophages (5.2%), and dendritic cells (4.2%) (**Fig. 4A, Fig. S5A-D, Table S5**). We further characterized these clusters of monocytes/macrophages based on comparison to publicly available gene expression data for monocyte clusters in the MoMac data set^32^, which integrated monocytes in health and disease states from 41 studies in 13 different tissues (**Fig. 4B, Fig. S6A, Table S6**). We also performed genes co-expression module detection and obtained 11 transcriptionally distinct modules (M1-M11), which helped us better define the cell states we observed (**Fig. 4C, Fig. S7A, Table S7**).

**Figure 4.**
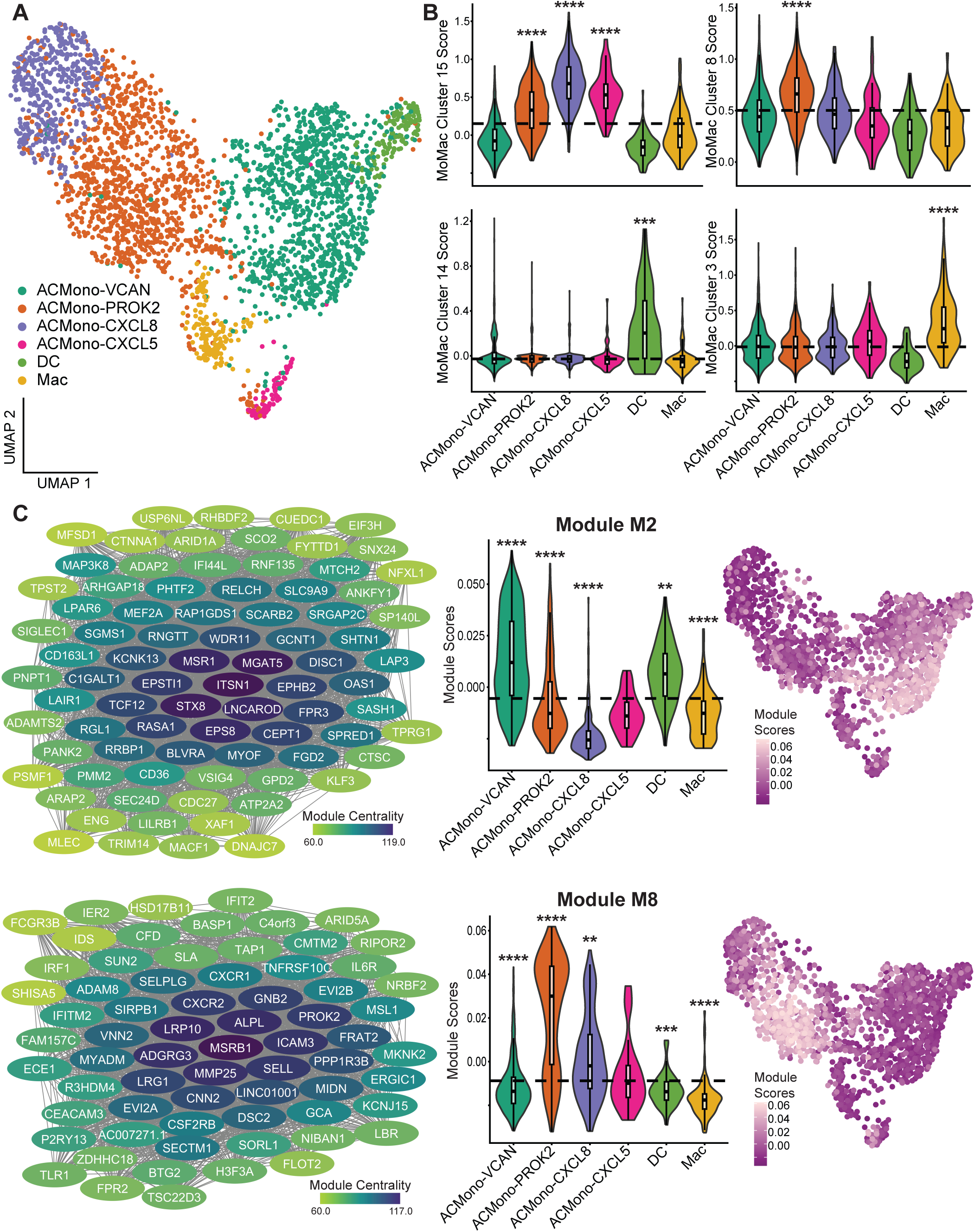
Monocyte diversity in CSF after IVH. A. UMAP indicating monocyte subtypes. B. Annotation of monocyte subtypes by scoring with marker genes from select subtypes identified in a meta-analysis of monocytes and macrophages from healthy and diseased samples (MoMac)^32^. C. Gene co-expression modules differentially expressed across monocyte subtypes. Wilcoxon rank-sum test (one-sided for reference marker scoring, two-sided for gene co-expression networks) with Bonferroni correction used for statistical significance testing (see Tables S6, S7). *p < 0.05, **p < 0.01, ***p < 0.001, ****p < 0.0001.

Classical monocyte markers (*SELL, CSF1R, ITGAX, CD163, CSF3R, S100A8*, *S100A9*, *S100A12, ALOX5AP*) were split between four distinct clusters that also expressed markers associated with monocyte activated or inflammatory states (*HLA-DRA, HLA-DRB1, CD163, IL1B, CXCL8, CCL3, CCL4, CXCL5,* members of the TNF family); hence we have labelled them Activated Classical Monocytes (AC Mono). We defined each of the four AC Mono subtypes by its most differentially expressed genes encoding secreted proteins: ACMono-VCAN (36.68% of all monocyte/macrophage cells), ACMono-PROK2 (36.1%), ACMono-CXCL8 (15.24%), and ACMono-CXCL5 (2.53%). The first two subtypes were the most abundant and had in common a pro-inflammatory signature including interferon-induced genes, while the latter two were defined by their high expression of neutrophil-attracting chemokines.

ACMono-VCAN was the most abundant monocyte subtype. Its eponymous marker *VCAN* (avg log_2_FC = 2.64, adjusted p = 1.53×10^-268^), encodes versican, a chondroitin sulfate proteoglycan that modulates inflammatory responses in many cell types^33,34^. It has been reported that *VCAN* activation in monocytes/macrophages is dependent on Type I interferon signaling^35^. Consistent with this, we observed elevated expression of the interferon-induced genes *IFI44* and *IFI44L.* ACMono-VCAN also had upregulation of genes in co-expression modules M2 (adjusted p = 1.35×10^-1^^18^) and M3 (adjusted p = 1.05×10^-11^^3^), enriched for viral defense responses and endocytic/phagocytic genes, respectively (**Table S7**). For instance, scavenger protein genes such as *STAB1* (avg log_2_FC = 1.83, adjusted p = 1.18×10^-117^) and *CD163* (avg log_2_FC = 1.76, adjusted p = 2.25×10^-83^) were upregulated, along with components of the THBS pathway (**Fig. 5C**), suggesting that this is an interferon-activated cell state responding through modification of the cell matrix. By contrast, ACMono-VCAN appeared not to be activated by other cytokines such as interleukin-1, as it had decreased expression of M5 (adjusted p = 7.42×10^-164^), a gene co-expression module enriched in genes involved in cellular response to interleukin 1 (GO:0071347), apoptotic process (GO:0006915), and NF-κB signaling (GO:0007249) module.

**Figure 5.**
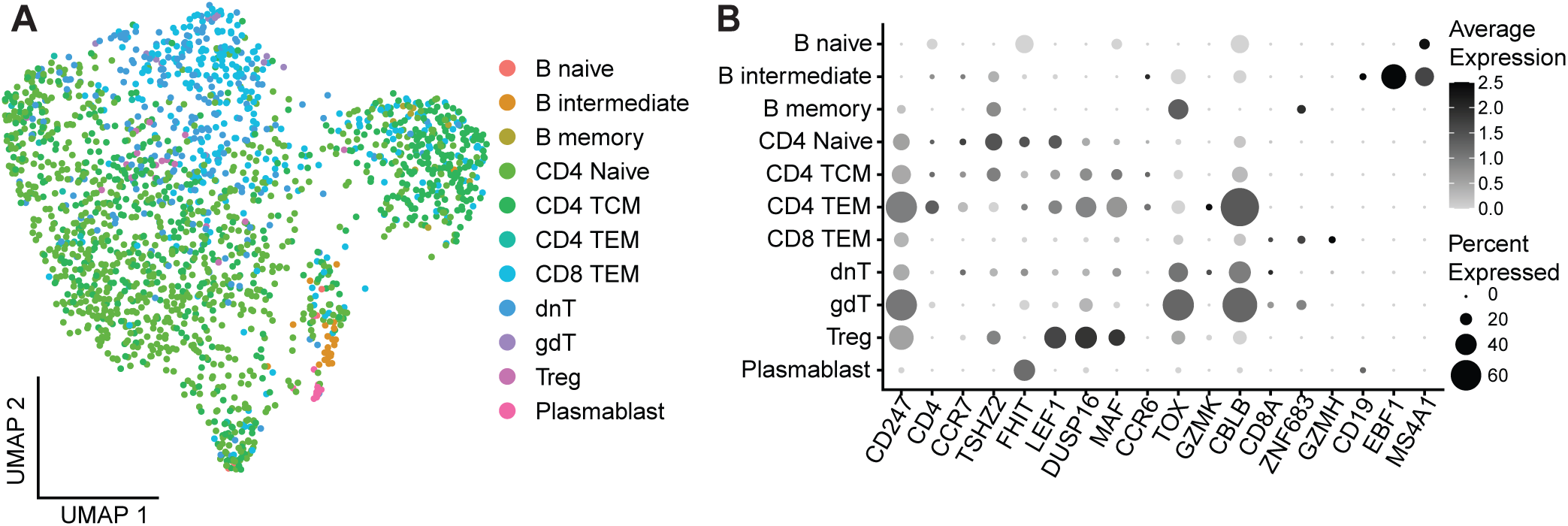
Lymphocyte diversity in the ventricular CSF. A. UMAP showing reference annotations in blood from the Azimuth data set^51^. B. Annotation of lymphocyte subtypes in CSF after IVH based on label transfer analysis. (C.,D.) Dot plots showing expression of marker genes in the reference set (C) and our lymphocyte population in CSF after IVH (D).

ACMono-PROK2 was characterized by the expression of *PROK2* (avg log_2_FC = 1.76, adjusted p = 1.68×10^-38^), encoding prokineticin-2, a peptide with chemokine–like activity that is highly expressed in many classical monocytes^36^ with roles in cell migration^37,38^ and differentiation^39,40^ and neurogenesis^41,42^. More broadly, the ACMono-PROK2 gene signature suggests a pro-inflammatory state, with elevated expression of gene co-expression module M8 (adjusted p = 3.52×10^-115^), including the cytokines *TNF*, *IL6*, and *IL1B*, and genes related to NF-κB signaling. Comparisons to MoMac revealed overlap with MoMac clusters 1 (adjusted p = 2.35×10^-126^) and 8 (adjusted p = 4.13×10^-101^), both of which are subtypes of CD16^+^ monocytes with high expression of inflammatory genes.

ACMono-CXCL8 was defined by elevated expression of pro-inflammatory ligands such as *CXCL8*, *CCL3*, and *CCL4*, markers of activated M1-like monocytes that act as chemoattractants for other monocytes, neutrophils, NK cells, and T-cells^43^. ACMono-CXCL8 had high expression of markers for Module M5 (adjusted p = 1.60×10^-142^), which was enriched for Gene Ontology terms related to vesicle secretion (<u>e.g.,</u> Proton-Transporting V-type ATPase (GO:0033180), positive regulation of exosomal secretion (GO:1903543), and multivesicular body secretion (GO:0032585) and with the interleukin-1-mediated signaling pathway (GO:0070498). Similarly, ACMono-CXCL8 had elevated expression of Module M9 (adjusted p = 6.07×10^-101^), which consists primarily of genes associated with vesicle production. This cluster also highly expressed markers of the “IL1B” MoMac Cluster 15 (adjusted p = 2.30×10^-141^), as well as the NF-κB transcription factor inhibitor *NFKBIA,* which regulates the pro-inflammatory activities of NF-κB downstream of TLR4. Collectively, these enrichments suggest that ACMono-CXCL8 may have M1-like activity.

ACMono-CXCL5, the smallest monocyte population, expressed activated monocyte ligands such as *CXCL5* and *CXCL3* that are known for their roles as chemoattractants for myeloid cells. In common with ACMono-CXCL8, ACMono-CXCL5 had elevated expression Module M9 (adjusted p = 2.85×10^-6^), and MoMac Cluster 15 (adjusted p = 2.95×10^-12^).

Macrophages (Mac) represented 5.24% of cells in the monocyte lineage. In addition to genes that are largely expressed in macrophages such as *GPNMB, PLD3* and *CTSD*, these cells were marked by high expression of pro-inflammatory genes such as *HIF1A-AS3*^44^ *FTL*^45^, and *MDM2*^46^. *FTL* expression in macrophages has been associated with exposure to oxidized low-density lipoprotein^47^. Consistent with this, we found high expression of several genes associated with lipid-associated macrophages (e.g., *CSTB*, *CTSD*, *GPNMB*, *LGALS3*, *PLD3*, and *PLIN2*^48^), which are also induced in macrophages in a murine model of stroke^49^. Interestingly, we also detected high expression for markers of MoMac cluster 3 (adjusted p = 1.50×10^-26^), a TREM2^+^, M2-like population. Overall, these annotations demonstrate the presence of macrophages in the CSF after IVH and provide evidence for heterogeneous reactive states.

The final sub-population (4.21%) expressed markers of dendritic cells (DC). We annotated these cells as DC based on shared markers with MoMac cluster 14 (adjusted p = 5.54×10^-22^), described by the MoMac authors as dendritic cell contamination. We note, however, that dendritic cells share several markers with intermediate monocytes, making a definitive annotation difficult. Some cells in the cluster expressed intermediate monocyte markers such as *CIITA*, suggesting a mixed population (**Fig. S5D)**. Notably, this population was also distinguished by elevated expression of MHC class II markers such as *HLA-DRA, HLA-DRB1, HLA-DQA1 and HLA-DQA2*, suggesting that these cells may have the ability to interact with and activate CD4⁺ T cells.

### Lymphocyte diversity in the intraventricular CSF

The 1,990 nuclei identified in our data as lymphocytes expressed canonical pan-lymphocyte and T cell markers, such as *PDE3B*, *FOXP1*, *MBNL1*, *ANK3*, *BCL2*, *FYN*, *SKAP1* and *CD247*. A small population of these cells expressed B-cell markers (*CD19*, *CD20*, *CD21*), consistent with observations for intraparenchymal hematoma and CSF in subarachnoid hemorrhage by flow cytometry^15,50^. *De novo* clustering of lymphocytes did not clearly define known T- or B-cell subtypes, most likely because of the relatively small number of cells representing each subtype. Therefore, to better characterize lymphocyte populations, we performed a label transfer analysis by integrating our lymphocytes with those from a large PBMC reference atlas and projected our data onto the UMAP from the reference data to identify shared marker genes and confirm annotations^51^ (**Fig. 5**). This analysis suggested that most of the lymphocytes we detected were CD4^+^ subtypes, especially central memory T-cells (28.5%) and CD4^+^ naïve T-cells (41.6%). CD8^+^ effector memory T-cells represented an additional 14.7% of lymphocytes. Double-negative CD4^-^CD8^-^ T-cells represented 10.4% of all lymphocytes, suggesting an enrichment of activate T-cell states. Consistent with this, further sub-clustering revealed a population expressing pro-inflammatory markers such as *S100A8*, *S100A9*, *CSF3R*, *IFITM2*, and *CXCR2*, a marker of γδ T-cells^52^.

### Intercellular communication networks in the intraventricular CSF

Chemokine ligands and their receptors were markers for several sub-populations of neutrophils, monocytes, and lymphocytes. For instance, the chemokine *CXCL8* was expressed almost exclusively by ACMono-CXCL8, while its receptor, CXCR2, was expressed primarily by neutrophils. The specific expression of these ligand-receptor pairs provides the potential for signaling between populations. To more rigorously investigate this putative cell-cell communication, we made use of Immune Dictionary^53^ and the CellChat^54,55^ package in R.

The Immune Dictionary provides single-cell transcriptomic profiles of 17 immune cell types in response to treatment with each of 86 cytokines in mice. We reasoned that similarities between cells in our data set and those responsive to specific cytokines in the Immune Dictionary imply common stimuli. We observed expression of IL-1α-induced genes in all three neutrophil populations (**Fig. 6A, Table S8**). In the IFN-activated neutrophil population, expression signatures of interferon responses (Type I and Type II) and IL-1 signaling (IL-1α, IL-1β, IL-15, IL-18, and IL-36α) were evident. While IFN-activated neutrophils were IL-1 responsive, they were not IL-1 productive, as they were negatively associated with co-expression module N10, a hub gene of which was *IL1B*. This is consistent with observations that Type I interferon can suppress IL-1 production^31^. In contrast, monocytes generally showed a greater expression of genes that are suppressed by IFN-α1 and IFN-γ stimulation (**Fig. S8 A-D**). Monocytes also expressed IL-1α- and IL-1β induced genes at high levels. Interestingly, PROK2 monocytes expressed higher levels of genes that were both induced and suppressed by IL-1α and IL-1β, suggesting a complex remodeling of the IL-1-modulated transcriptome. ACMono-VCAN, ACMono-CXCL5, and DC cells expressed a signature observed in monocytes treated with IL-33 (**Fig. 6B**), which is an IL-1 family cytokine that is released in response to cell stress or damage. Macrophages shared with monocytes an IFN-α1 and IFN-γ suppressed expression profile, but also highly expressed genes that were suppressed by IL-1α.

**Figure 6.**
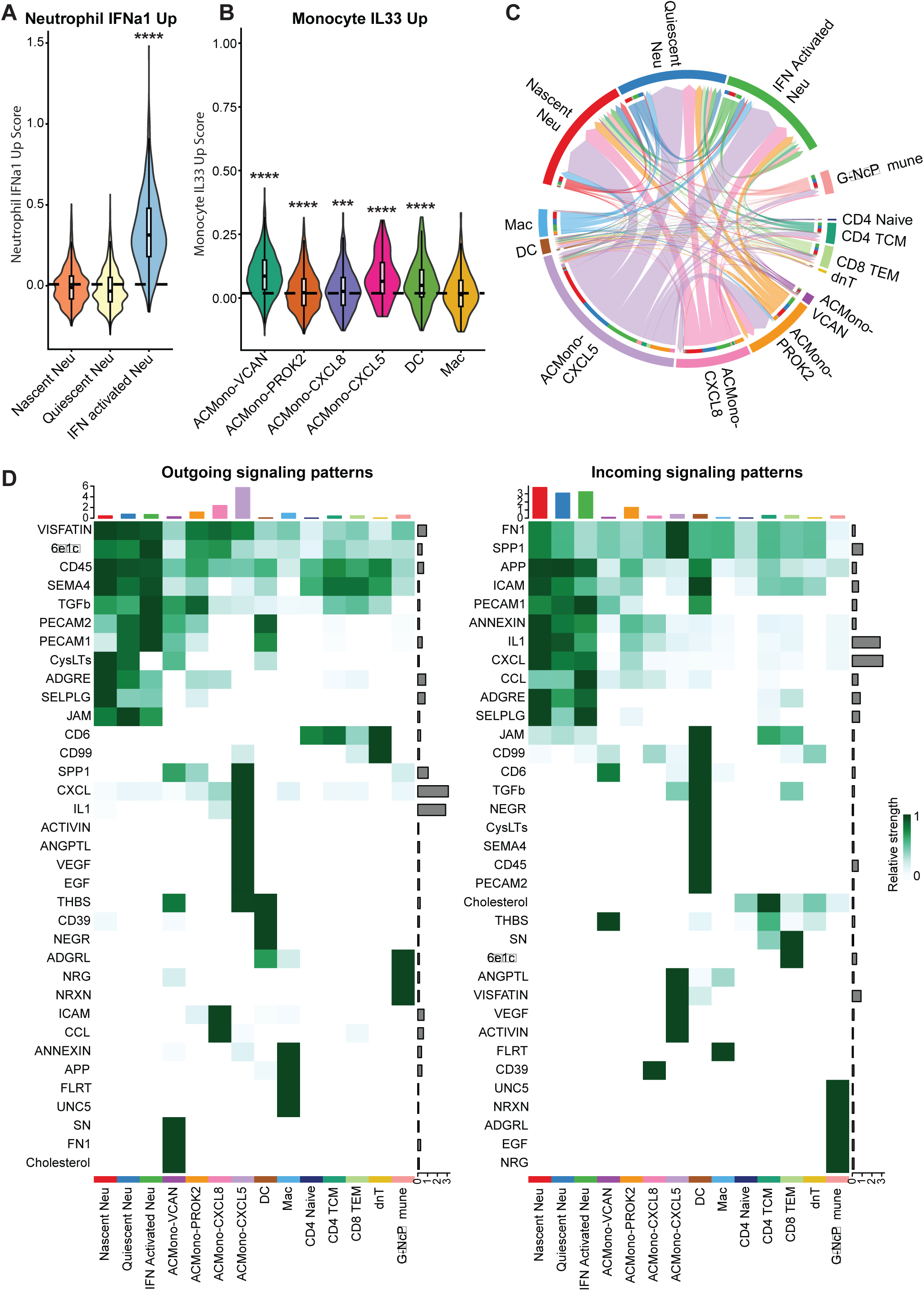
Intracellular communication in ventricular CSF. A. Neutrophils in CSF scored by genes upregulated in Neutrophils from the Immune Dictionary upon IFNa1 exposure. B. Monocytes, dendritic cells and macrophages in CSF scored by genes upregulated in Monocytes from the Immune Dictionary upon IL33 exposure. C. Circos plot showing strength of outgoing and incoming signals across cell states in the vCSF, based on CellChat analysis. D. Heatmap of outgoing and incoming signal patterns for cytokine classes across all the cell states in the vCSF, based on CellChat analysis.

**Figure 7.**
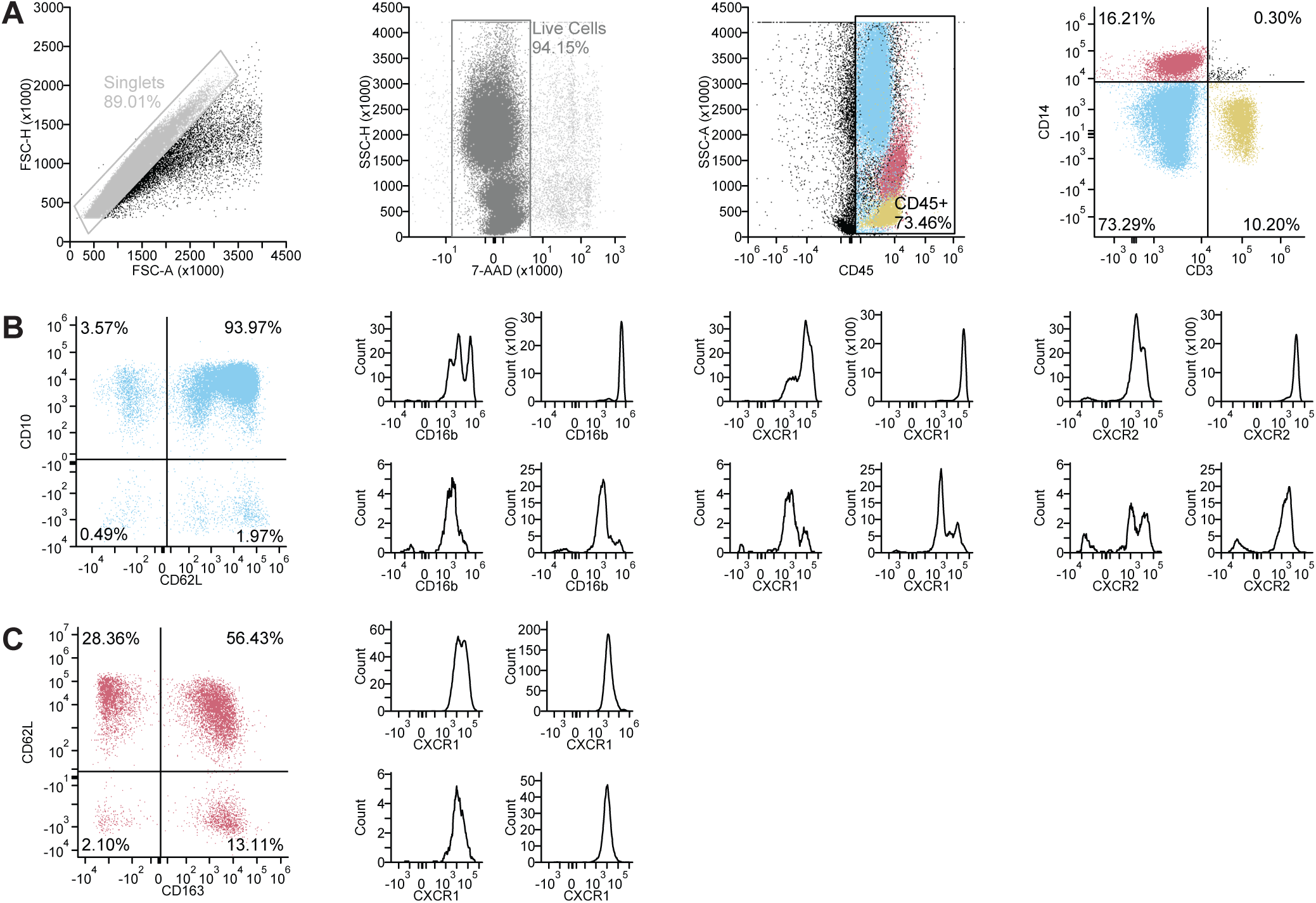
Flow cytometry of CSF after IVH. Aggregate data from n=3 samples from n=2 patients is shown, for a total of 72,725 events. A. Gating strategy for identifying the singlet, live, CD45^+^ cell population (53,424 events) with CD3 and CD14 used to distinguish neutrophils (orange), monocytes (blue), and lymphocytes (green). Back gating onto the SSC-A vs. CD45 plot shows separation of these populations based on the characteristic distribution of these cells types by side scatter. B. Neutrophil population is further gated by CD10 and CD62L. Each quadrant is subsequently gated by either CD16b, CXCR1, or CXCR2. C. Monocyte population is further gated by CD62L and CD163. Each quadrant is subsequently gated by CXCR1.

We further delineated intercellular communication networks using CellChat, which uses the expression patterns of receptor-ligand pairs to identify cell types with the potential for cell-cell communication. Doing so, we found that the expression of ELR-motif CXC chemokines linked ACMono-CXCL5 and ACMono-CXCL8 to chemokine receptor CXCR1- and CXCR2-expressing neutrophils. This pathway is particularly important for neutrophil recruitment and activation, suggesting a role for macrophages in facilitating this function. In addition, ACMono-CXCL8 contribute CC chemokines that tie them to IFN-activated neutrophils specifically. CellChat also revealed interactions between subtypes of neutrophils, mediated by ADGRE and SELPLG/SELL. Other interactions are shown in **Fig. 6C and 6D**.

Integrating observations from Immune Dictionary and CellChat suggested complex behaviors not evident from either data set alone. For example, from Immune Dictionary we observed that ACMono-CXCL5 shared common gene expression features with macrophages stimulated by the IL-1 family cytokine IL-33. CellChat indicated that ACMono-CXCL5 also contribute IL-1, suggesting an IL-1 signaling positive feedback loop in this macrophage subtype.

### Development of a flow cytometry panel for neutrophil and monocyte diversity in the ventricular CSF

Our snRNA-seq data suggest that specific immune cell subtypes are enriched in the CSF of IVH patients compared to their prevalence in other contexts. Monitoring the abundance and dynamics of these immune cell subtypes could potentially reveal biomarkers that predict clinical outcomes. However, snRNA-seq is not a practical method to track these populations in clinical settings. Leveraging our snRNA-seq data, we sought to develop a panel of cell surface markers to distinguish neutrophil and monocyte subtypes by flow cytometry.

We therefore developed a flow cytometry panel to delineate functional states of neutrophils and monocytes, the two most common cell types observed in our snRNA-seq data. To leverage the insights of our snRNA-seq data, we derived this panel empirically from our data set, as opposed to relying on existing cytometry-based classifications. After identifying cluster of differentiation (CD) genes enriched in specific clusters, we used FluoroFinder (Broomfield, CO) to select a subset of markers for which reliable antibodies with compatible fluorophores existed.

Using this panel, we performed flow cytometry analysis of CSF samples on three samples from two participants with IVH not included in the snRNA-seq analysis. Both participants had experienced thalamic ICH with IVH, one as a result of hypertension and other as a result of a ruptured arteriovenous malformation. As expected, CD45+ immune cells could be distinguished as granulocyte, monocyte, and lymphocyte populations using CD3 and CD14 gating, which were further confirmed by forward and side scatter profiles (**Figure 6A**). Based on neutrophil clustering from our snRNA-seq data, we hypothesized that three canonical neutrophil surface markers (CD10, CD62L, and CD16b) could help resolve distinct neutrophil subsets^56^. The dominant population (>90% of neutrophils) was CD10^+^CD62L^+^CD16b^high^, consistent with mature circulating neutrophils.

CD10 presented a particular interpretive challenge: while transcriptional data, including our own and that from Wigerblad et al.^16^, suggest CD10 as a marker of nascent neutrophils, it is better established as a surface protein associated with mature neutrophils. Consistent with this, we observed a small population of CD10^-^CD62L^+^CD16b^high^ neutrophils (<2%) consistent with a nascent phenotype, but no CD10^+^CD62L^+^CD16b^high^, as otherwise might have been predicted directly from our transcriptomic data **(Figure 6B)**. From our transcriptomic data, we would infer that quiescent neutrophils are CD10^-^CD62L^-^CD16b^low^. However, we again anticipated that the protein-level expression of CD10 would differ for these cells, and that quiescent cells should be CD10^+^CD62L^-^CD16b^low^. Though both populations are small, the CD10^+^CD62L^-^ CD16b^low^ population is much more common in our flow cytometry data than is the CD10^-^CD62L^-^ CD16b^low^ population **(Figure 6B).** A distinct subset of a CD10^-^CD62L^-^ population is CD10^-^CD62L^-^ CD16b^high^. Cross-referencing with our snRNA-seq data, this population is most consistent with interferon-activated neutrophils, though the markers shown here do not directly test for that state. Additionally, we identified by flow cytometry a CD10^+^CD62L^-^CD16b^+^ subset not resolved in our snRNA-seq clustering analysis but reported in the literature as T cell–suppressive neutrophils^57^. Similarly, we hypothesized that monocyte subtypes could be distinguished using the markers CD163, CD62L, CXCR1, CXCR2, CD50, and CD87. We found that of these, CD163, CD62L, and CXCR1 were the only markers with discriminating power in our flow cytometry data set **(Figure 6C**).

We note that we detected nascent, quiescent, and interferon-activated populations — which together dominate the snRNA-seq neutrophil profile -- at very low levels in our flow cytometry data. It is unclear whether this arises from greater sensitivity of snRNA-seq; misclassification by snRNA-seq due to mismatch between gene and protein expression; limitations of the flow cytometry panel; or patient-specific differences in neutrophil subtype. Taken together, however, our flow cytometry results indicate that it may be possible to translate leukocyte functional states inferred from single-nucleus transcriptomics into clinically-feasible flow cytometry panels.

## Discussion

In this study, we generated a single-nucleus transcriptomic atlas of the ventricular CSF in IVH. We leveraged nuclear extraction as a strategy to enrich for cell types — specifically neutrophils — that are often challenging to capture in single-cell experiments because of their fragility. Our results revealed a diversity of cell types, including distinct subpopulations of neutrophils, monocytes, and lymphocytes. Notably, we captured a large and diverse population of neutrophils, including a population of interferon-activated neutrophils that has not been described in the central nervous system previously. As single-nucleus RNA sequencing is not feasible for clinical diagnostic purposes, we validated potential flow cytometry markers for cellular sub-clusters we identified. This demonstrates the feasibility of translating biomarkers derived from snRNA-seq data into clinically practical flow cytometry markers.

A strength of our study that distinguishes it from previous studies is the detection of a large population of neutrophils in the CSF. Neutrophils are expected to be among the most common leukocytes present in the CSF during the first several days after hemorrhage^58^. Neutrophil depletion has been shown to be protective in a range of hemorrhagic brain disease models^59–61^. Yet previous sequencing studies in these diseases have selected against neutrophils in sample processing. Sansing and colleagues reported single-cell RNA sequencing data from the intraparenchymal hematoma of a patient who underwent clot evacuation after an intracerebral hemorrhage as part of the MISTIE III phase III clinical trial^62^ and have recently extended this work to a larger group of subjects undergoing clot evacuation^63^. Their study provides a rich time-dependent atlas of leukocytes in the intracerebral hematoma, but only a small population of neutrophils was detected because they subjected samples to a specific granulocyte depletion. Similarly, the two reported single-cell studies of CSF in subarachnoid hemorrhage^64,65^ specifically selected against neutrophils, one profiling PBMCs and the other profiling CD8^+^CD161^+^ cells. Until recently, neutrophils were thought to display a relatively limited functional repertoire, owing to their relatively low transcriptional activity and short lifespans. Single-cell RNA sequencing has revealed that, in fact, neutrophils are highly diverse cell populations^16,66,67,68^. This expanded understanding of the diversity of neutrophils prompted us to investigate this cell population more comprehensively in IVH.

Neutrophils may be particularly challenging to capture in single-cell studies. They are often short-lived and highly sensitive, rendering them difficult to capture with methods that have long or complex workflows. This raises the question of why we were able to identify a large population of neutrophils when others have not. We analyzed CSF instead of parenchymal hematoma, and these may have different cell populations. However, numerous neutrophils can be isolated from parenchymal hematoma by flow cytometry^69^. It is likely that differences in our sample preparation account for the preservation of neutrophils. The use of single-nucleus instead of single-cell RNA sequencing may improve the identification of neutrophils, as all cells undergo lysis, reducing the bias against fragile neutrophils. While we cannot fully exclude the possibility that the nuclei we identified as deriving from neutrophils actually arise from other cell types, the expression of canonical neutrophil markers in these cells supports their assignment.

A strength of single-nucleus RNA sequencing over traditional cytometric approaches is the ability to distinguish subtle cellular phenotypes and activation states. The advent of these methods has revealed that neutrophils, previously thought to be a relatively homogenous cell type, in actuality exhibit a variety of states^16^. In this case, we identified three major neutrophil functional states, though we anticipate that with larger sample sizes and improved isolation techniques, it is likely we would be able to classify these states even more specifically. We have shown that these are consistent with classifications of neutrophils derived from peripheral blood in independent data sets^16,19^.

One particularly interesting neutrophil state is the IFN-activated neutrophil. To our knowledge, this subtype of neutrophils has not been reported in the human central nervous system previously, though it has been detected in rodents^70^. Interferons have been known to stimulate neutrophils for nearly half a century^71^. It was recently reported that interferon-activated neutrophils play an important role in tumor control^17,72^ and the response to SARS-CoV-2 infection^18^. Knockout of the Type I interferon receptor results in increased neutrophil accumulation in infected tissue^73,74^, while interferon-gamma decreases neutrophil accumulation in such tissue^75^. Our analysis based on Immune Dictionary suggests that Type I and II responses may be particularly relevant to IFN-activated neutrophils in IVH.

A study of soluble factors in the CSF after IVH identified time-dependent increases in the Type II interferon IFN-γ that were augmented in patients treated with intraventricular tissue plasminogen activator; no subjects in our study underwent such treatment^76^. Similarly, IFN-γ levels in the CSF were associated with white matter damage in premature infants with posthemorrhagic hydrocephalus^77^. Little is known about the role of Type I interferons in human IVH, though a nonsense single-nucleotide polymorphism in the gene encoding IFN-ε is associated with the development of ICH^78^. In a rat germinal matrix hemorrhage model, treatment with Rh-IFN-α improved neurological function^79^. Our findings suggest that the Type I interferon response in general, and interferon-activated neutrophils in specific, merit further investigation in the pathophysiology of intraventricular hemorrhage. This represents a potentially targetable pathway for modulating neutrophils for future therapies.

Our data also reflect a diversity of monocytes/macrophages that occupy a range of functional states. These states corresponded to strong differential expression of genes encoding certain secreted proteins, which may give some indication to their role in IVH and other acute brain injuries. The distinction we observed between macrophages expressing lipid metabolism genes and monocytes expressing cytokine-stimulated genes was previously noted for tumor-associated cells^32^. We further classified monocytes into four activated states defined by their expression of genes encoding specific secretory proteins. *VCAN ACMono* and *PROK2 ACMono* exhibited transcriptomic profiles that could be described broadly as pro-inflammatory and interferon-responsive, gene co-expression network analysis revealed markedly different profiles for these cellular states. Increased deposition of versican (encoded by *VCAN*), a protein that is involved in myelination, has been noted in the brains of rabbit pups subjected to IVH^80^. Overexpression of *PROK2* has been shown to reduce neurodegeneration and functional deficits in a mouse model of TBI^81^. Both the VCAN ACMono and PROK2 ACMono express interferon-response genes, again supporting an outsize role of interferon signaling in the immune response to IVH. Further insight can be gleaned receptor-ligand interactions, such as CXCL8 and CXCR2, which we have characterized more extensively using CellChat. These results indicate cross-talk within subtypes of individual cell types (e.g., PROK2 with CXCL8 ACMono) or between cell types (e.g., PROK2 ACMono with IFN-activated neutrophils). Major roles for the ELR-motif CXC chemokines, CC chemokines, IL-1 family cytokines, and interferons were observed. We also observed monocyte/macrophage phenotypes that have been associated with disease. For example, both snRNA-seq and flow cytometry revealed a substantial number of CD163^+^ monocytes/macrophages, a population that has been associated with disease severity in aneurysmal subarachnoid hemorrhage^82^.

The distribution of lymphocytes we observed in the CSF is intriguing and merits further investigation. The majority of cells were CD4^+^ T cells. It has been reported that the majority of T cells in the CSF under normal conditions exhibit a central memory phenotype^83^. This was one of the major subtypes we observed in IVH, though not the most common. These cells have been purported to enter the CSF via the choroid plexus^84^.

Limitations of our study include a small sample size and insufficient material for genotyping, both of which reduce our ability to delineate differences inflammatory profiles that are associated with disease severity. These samples were convenience samples that were obtained when specific clinical events (EVD placement, EVD removal, or diagnostic testing) prompted CSF sampling, increasing the heterogeneity of the samples and potentially creating sampling bias. In addition, low cell counts from individual subjects further limit the generalizability of our results. Nevertheless, our study is one of very few single-cell or single-nucleus transcriptomic studies of cerebrospinal fluid for hemorrhagic brain disease, and substantially increases the data available in the literature. While we hypothesize that the single-nucleus approach has allowed us to capture difficult-to-detect neutrophil states, this approach does have drawbacks. Nuclei contain substantially fewer transcripts than do whole cells, potentially diminishing our ability to identify transcriptionally regulated pathways relevant to disease biology. Nevertheless, even small nuclear transcriptomes have proven reliable for cell type classification in our study. While single-cell transcriptomic methods are unlikely to be translatable directly to clinical diagnostics, they may generate candidates for the development of flow cytometry panels. We provide a proof-of-concept for this here, but with too few subjects to demonstrate the utility of these markers for diagnostic or prognostic purposes.

It would be valuable to extend our approaches to elucidate time-dependent changes in blood and CSF leukocytes across a larger cohort of patients in ICH, SAH, IVH, and TBI. Previous studies report divergence in leukocytes sampled from the blood and central nervous system over time^15,62^. The distinct immune profile in the CSF and brain after IVH may yield insights into the risk for common complications, such as persistent hydrocephalus. Such information could help to inform the timing of surgical interventions and reveal strategies to guide treatment of post-hemorrhagic hydrocephalus.

Taken together, our results provide the most comprehensive cellular atlas of the CSF in IVH to date. This atlas reveals a rich diversity of cellular populations, including myeloid states not previously described in the human central nervous system, including interferon-responsive neutrophils. We have further demonstrated that transcriptional markers from these populations can in many cases be validated at a protein level with flow cytometry. We anticipate that these results will set the stage for larger studies that elucidate the relevance of specific leukocyte subtypes to disease severity, progression, and outcome. Eventually, this will yield new diagnostic tools and therapeutic targets for intraventricular hemorrhage, a common, catastrophic disease with few effective treatments.

## Materials and Methods

### Sex as a biological variable

As we collected convenience samples and batched nuclei for sequencing (see below), we were unable to interrogate sex as a biological variable specifically; N=6 samples were from male subjects and N=1 sample was from a female subject. For the period of enrollment, a recruitment flow chart is shown in **Figure S9**.

### Statistics

Statistical methods used in this study are described in the sections pertaining to each specific assay or analytical tool.

### Study approval

Subjects (N=7; ICH=6, SAH=1) are a subset of those included in a prospective cohort study of patients aged 18-88 with potentially life-threatening neurologic illness (*NCT04189471*). The University of Maryland, Baltimore, institutional review board (IRB) approved this study, entitled “Improving Outcomes for Patients with Life-Threatening Neurologic Illnesses,” approval: number HP-00056063, date: August 1, 2013. Written informed consent was obtained from subjects (if they had decision-making capacity) or their legally authorized representatives. Study procedures were performed in accordance with the ethical standards of the responsible committee on human experimentation and with the Helsinki Declaration of 1975. Detailed clinical characteristics of study participants were recorded in a secure database, including Cytospin cytology and computed tomography (CT) scans obtained from the participants as part of the standard of care.

### Sample preparation and single-nucleus RNA sequencing

Of the CSF aspirated from the proximal port of the EVD during routine clinical practice, excess sample was obtained and transported to the laboratory for immediate processing. Obtained CSF was collected on EDTA (purple top vacutainers), with a processing time <1 hour from the time of collection. RNAse and Proteinase/phosphatase were inhibited by adding 1uM DEPC (Sigma D5758) and 1XHALT (Invitrogen 1861281). The cellular component of CSF was obtained by spinning the sample at 350g for 10 min. The pellet was resuspended, and red blood cells were depleted using EasySep magnetic beads (StemCell Technologies cat18170), and Miltenyi microcolumns (Miltenyi cat130-042-701). Samples were then washed and stored at - 80C. Half of the samples were stored as pellets and half in cryopreservation media (Cryostor, StemCell Technologies cat 100-1061). All the samples were stored at -80°C for <3 months. For pelleted samples, nuclei extraction was performed directly from the pellets. For samples stored in cryopreservation media, nuclei isolation was performed on the cellular fraction after washing with PBS+1% BSA at 350g/5 minutes at 4C. Nuclei isolation was performed using a detergent-free nuclei isolation kit, Invent (cat IN-024), according to the manufacturer’s instructions. Briefly, nuclei were resuspended in buffer A, then vortexed for 20s at level 9 on Genie2 Vortex, then placed on the spin column at 16000g/30s. The pass-through was washed with buffer B and counted on a MoxiGoII (Orflo) using 1uM Propidium Iodide (Sigma P-4864). We loaded two pools, one pool containing samples stored as pellets and other with cellular fraction cryoprotected. A total of 30K isolated nuclei per pool in PBS+1% BSA were loaded in a 10X Chromium X. Sequencing libraries were prepared by the Maryland Genomics core facility at the Institute for Genome Sciences using Chromium Single Cell 3’ Kit v3.1 reagents, according to manufacturer’s instructions. Libraries were sequenced on a NovaSeq6000 sequencer (Illumina) to an average depth of at least 25,000 raw reads per cell.

### Flow cytometry

CSF samples were acquired from the EVD buretrol chamber. Samples were stabilized in EDTA tubes during transport from the intensive care unit to the laboratory and kept at room temperature. Samples were then spun at 300g for 5 min, at 4°C. Pellets were then resuspended in 100 µL of staining buffer (phosphate buffered saline with 1% bovine serum albumin and 1% EDTA - A9418 and 324506, Sigma Aldrich), and incubated for ten minutes on ice with Fc Block (ThermoFisher). Pellets were resuspended in 50 µL of staining buffer mixed with antibodies and incubated for an additional 15 minutes in dark. A 13-antibody panel was used for this experiment. Antibodies included mouse anti-human anti-CD45 (363-0459-41, ThermoFisher), mouse anti-human anti-CD3 (344825, BioLegend), mouse anti-human anti-CD14 (367109, BioLegend), human anti-human anti-CD16b (130-123-643, Miltenyi Biotec), mouse anti-human anti-CD163 (333611, BioLegend), human anti-human anti-CD50 (130-115-309, Miltenyi Biotec), mouse anti-human anti-CD183 (353721, BioLegend), goat anti-human anti-CD121a (FAB269F-025, R&D Systems), mouse anti-human anti-CD62L (567989, BD Biosciences), mouse anti-human anti-CD10 (312241, BioLegend), human anti-human anti-CD181 (130-115-950, Miltenyi Biotec), human anti-human anti-CD182 (130-131-233, Miltenyi Biotec), mouse anti-human anti-CD87 (749063, BD Biosciences). 100 µL of staining buffer were added, cells were mixed and then spun at 300g for five minutes at 4° C. The supernatant was removed, cells resuspended in 200 µL of staining buffer, and 2 µL of 7-amino-actinomycin D (7-AAD) (420404, BioLegend) were added before flow acquisition. Flow acquisition was performed on a Cytek Aurora 4 Spectral Cytometer (Cytek Biosciences, Freemont, CA, USA) (lasers: 355nm, 405nm, 488nm, 640nm) at the University of Maryland Greenebaum Comprehensive Cancer Center Flow Cytometry Shared Service. At least 5,000 events were acquired and analyzed with FCS Express V7 software (De Novo Software, Pasadena, CA, USA).

### snRNA-seq cell clustering and annotation

Initial data processing was performed with CellRanger (7.2.0), yielding a cell x gene matrix for each pooled sample. Seurat^51^ v4.0.5 was used for batch integration and clustering, using the following parameters. We selected high-quality cells with at least 300 unique molecules and a maximum of 20 percent mitochondrial counts. SCT integration was performed with 5000 variable genes and 20 anchors. Louvain clustering was performed using 20 principal components. An optimal clustering resolution was determined by using the mrtree clustering optimization method^85^, which was given resolutions from 0 to 5, with 3.5 (35 clusters) being optimal for the entire population. Cell clusters were annotated to established cell types based on canonical markers from immune cells in other contexts^86–88^. Subsequently, the most abundant major cell types — neutrophils and monocytes — were sub-clustered, following the same procedures starting from the SCT integration step. Cellular populations derived from sub-clustering were compared to neutrophil and monocyte data sets that have been described by scRNA-seq in other contexts^16,19,32^. Cells in our snRNA-seq dataset were scored using the combined expression of marker genes from comparison datasets with the AddModuleScore() function in Seurat. Genes from reference datasets were selected with a log_2_FC > 0.585 (greater than 1.5 fold change) and adjusted p-value < 0.05. We also considered cell annotations derived from label transfer using the Seurat FindTransferAnchors and TransferData functions with 30 principal components and otherwise default parameters. Label transfer was performed separately for monocytes and lymphocytes using level 2 annotations from scRNA-seq of human peripheral blood mononuclear cells^51^.

### Gene co-expression networks in neutrophils and monocytes

Gene co-expression modules were identified through k-means clustering of imputed read counts, as previously described^89^. First, we performed a zero-preserving imputation with ALRA^90^ to reduce dropouts and improve correlation structure. Next, we performed k-means clustering (parameters). Modules with tightly correlated centroids were merged, and genes with insufficient correlation to the centroids of the merged modules were dropped. These steps were performed using the cells from Pool 2, then projected onto the cells from Pool 1. This strategy reduces the potential for batch effects and provided an independent test set to evaluate the robustness of the modules. Gene Ontology annotations for the modules were obtained using the enrichR package in R from the GO Molecular Function 2023, GO Cellular Component 2023 and GO Biological Process 2023 databases.

### Cell-cell signaling networks

Cytokine signaling activity among cell types was predicted by two methods: We used the CellChat^54^ (v 2.2.0) package in R to predict the incoming and outgoing signals and the associated pathways for each cell type. Cell populations with fewer than 50 cells were excluded due to insufficient data. Incoming) cytokine responses were further evaluated by scoring each cell (for the activity of signatures defined by the set of genes up- or down-regulated (FDR < 0.05) in immune cell populations after exposure to each of 86 cytokines^53^. As the Immune Dictionary was generated in mice, our analysis used human genes with one-to-one orthology. Scores were calculated with Seurat AddModuleScore() for each cytokine / cell type condition in which at least ten differentially expressed genes were detected.

## Data availability

We have made our expression data available on the Neuroscience Multi-Omic Analytics (NeMO) platform^91^ (https://nemoanalytics.org/p?s=c4d7a449) to facilitate secondary data visualization and analysis. Sequencing data will be available on dbGaP (pending). Code generated for this study is available at https://github.com/Ciryam-Lab/vCSF-Hemorrhage.git.

## Figure Legends

**Supplementary Figure S1.**
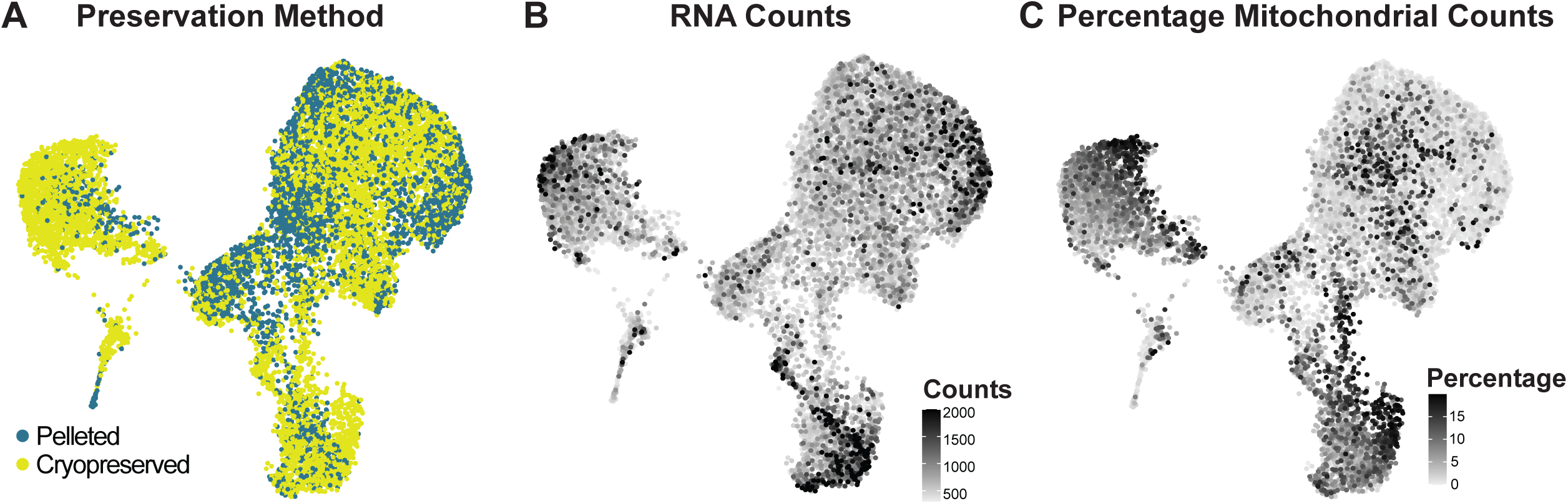
UMAPs of ventricular CSF colored by quality metrics. UMAPs of snRNA-seq of ventricular CSF (11,191 nuclei from seven study participants) colored by A. Preservation method. B. RNA Counts. C. Percentage Mitochondrial Counts.

**Supplementary Figure S2.**
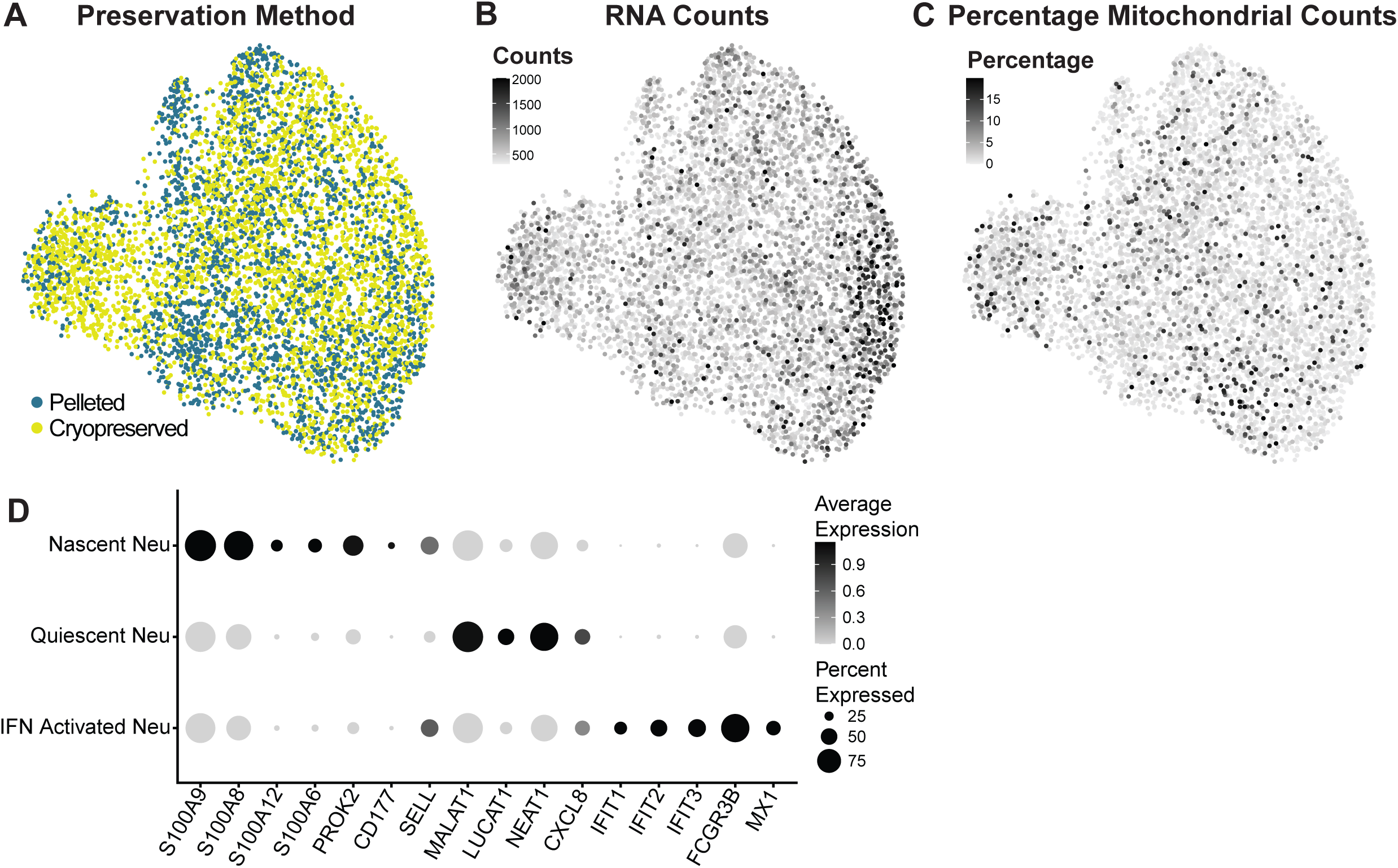
UMAPs of the neutrophil subset colored by quality metrics. UMAPs of snRNA-seq of the neutrophil subset colored by A. Preservation method. B. RNA Counts. C. Percentage Mitochondrial Counts. D. Dot plot showing marker genes for neutrophil subtypes.

**Supplementary Figure S3.**
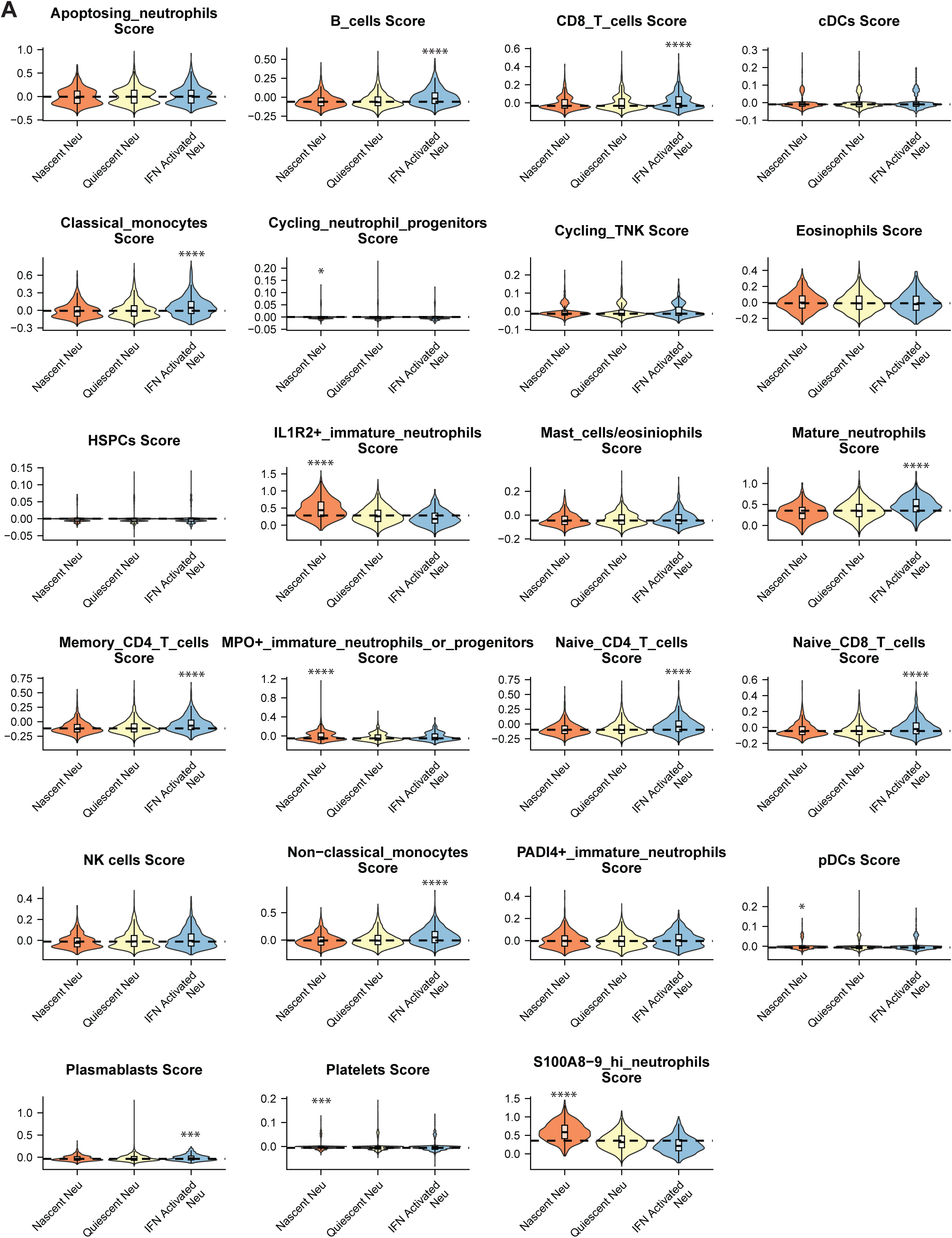
Annotation of neutrophil subtypes. Violin plots of neutrophil subtypes scored by marker genes from cell states found in blood during sepsis response^19^.

**Supplementary Figure S4.**
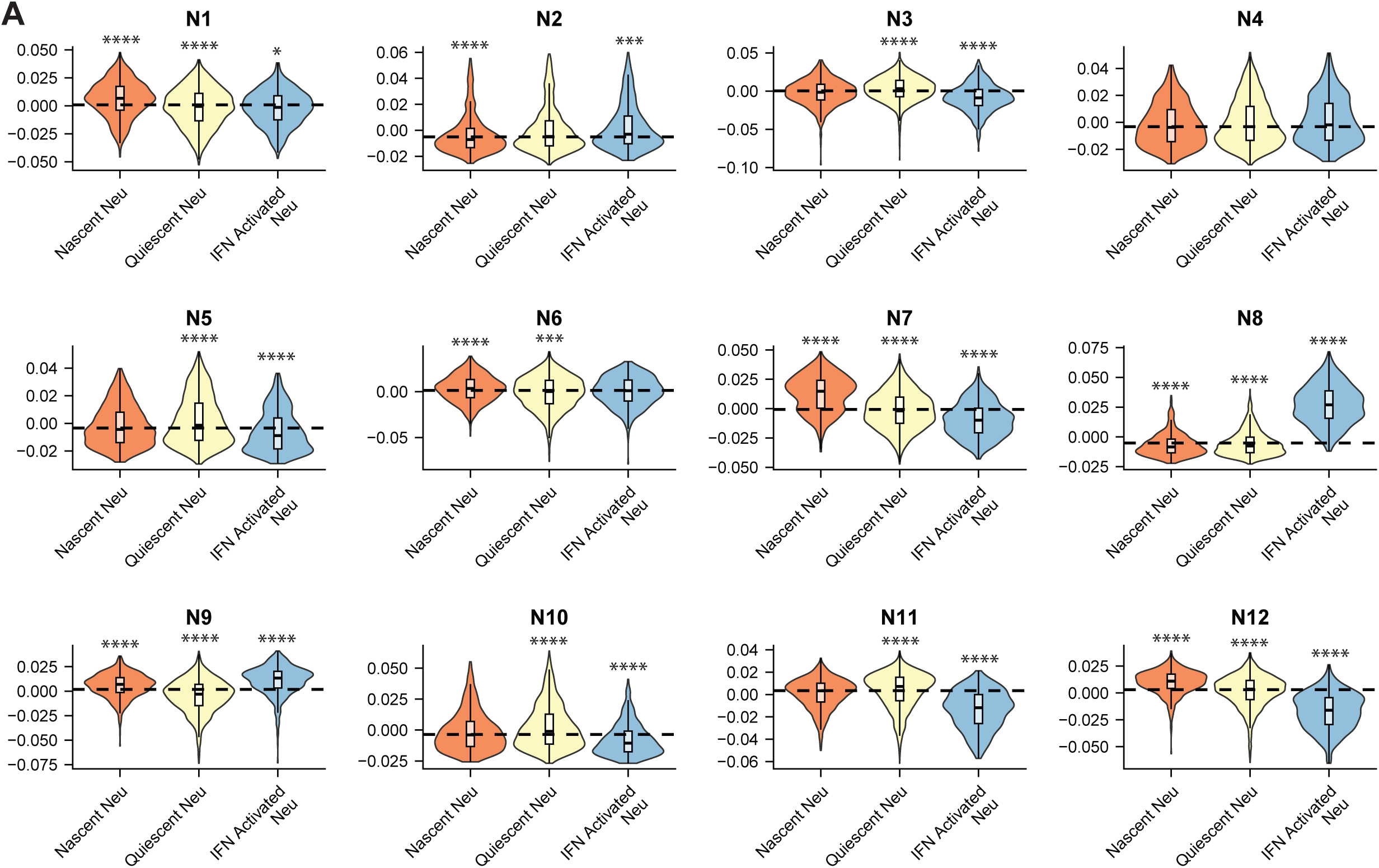
Gene co-expression Modules. 12 neutrophil gene co-expression modules differentially expressed across neutrophil subtypes.

**Supplementary Figure S5.**
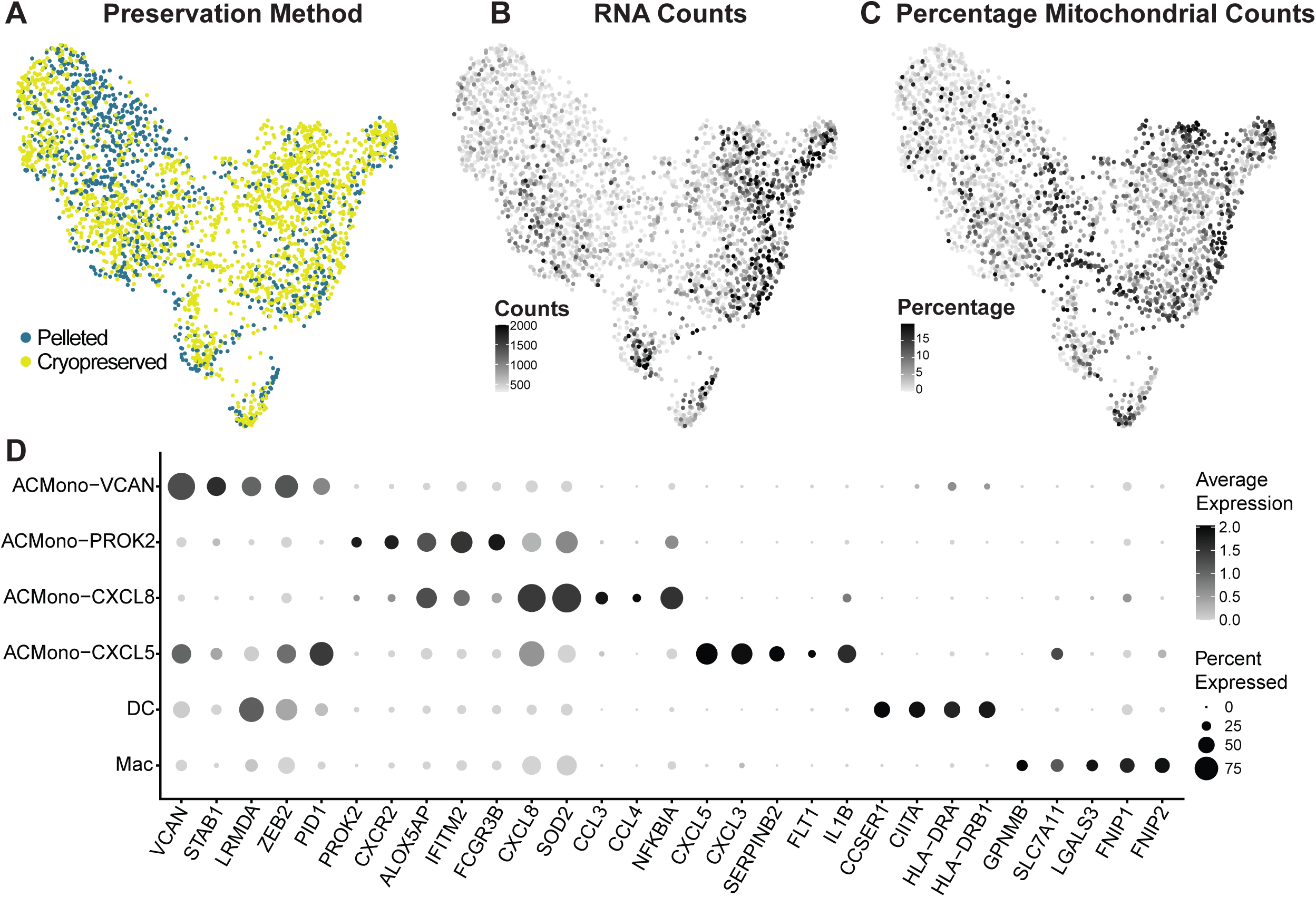
UMAPs of the monocyte subset colored by quality metrics. UMAPs of snRNA-seq of the monocyte subset colored by A. Preservation method. B. RNA Counts. C. Percentage Mitochondrial Counts. D. Dot plot showing marker genes for monocyte subtypes.

**Supplementary Figure S6.**
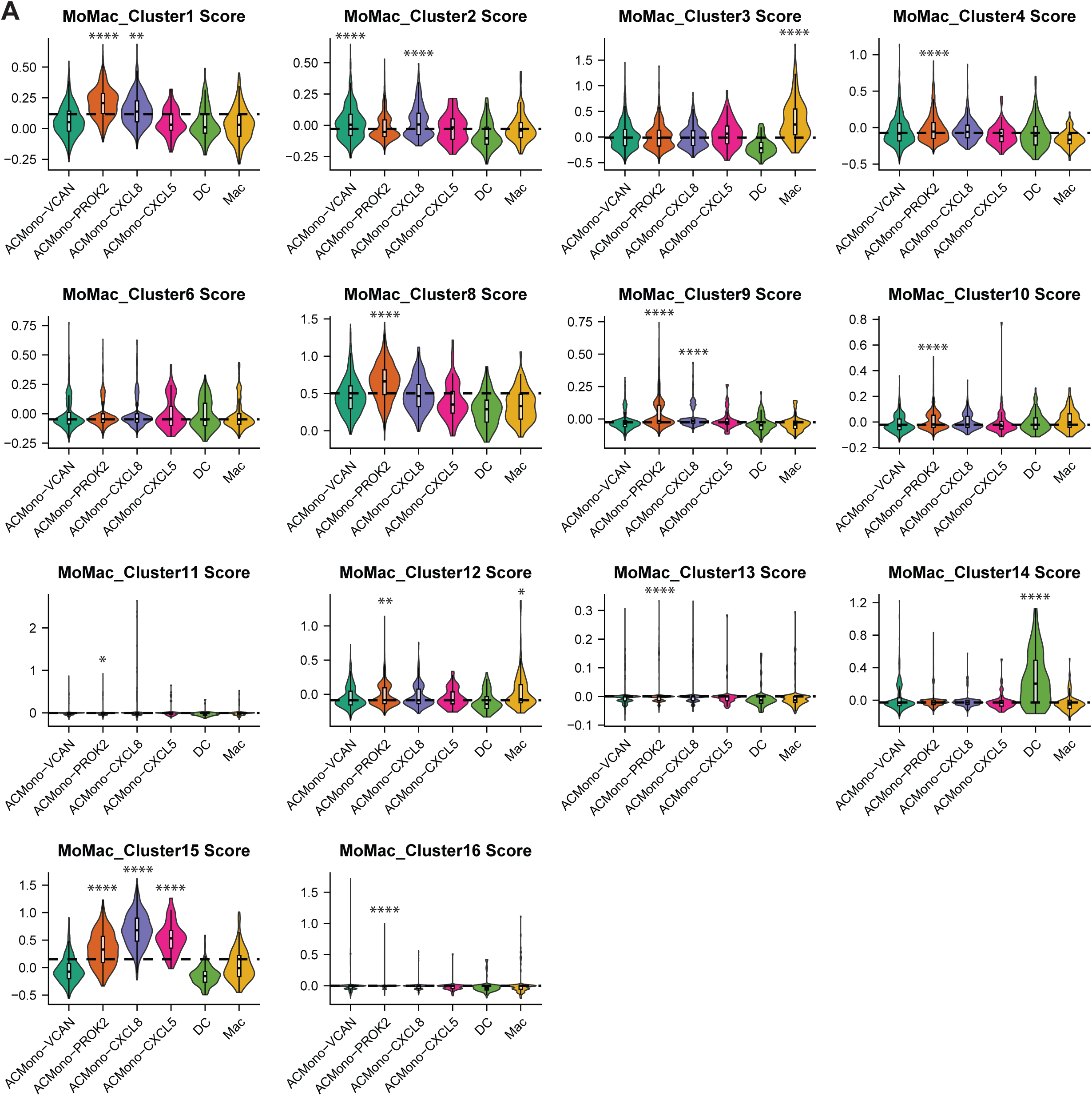
Annotation of monocyte subtypes. Violin plots of monocyte subtypes scored by marker genes from all the cell states found in the MoMac dataset^32^.

**Supplementary Figure S7.**
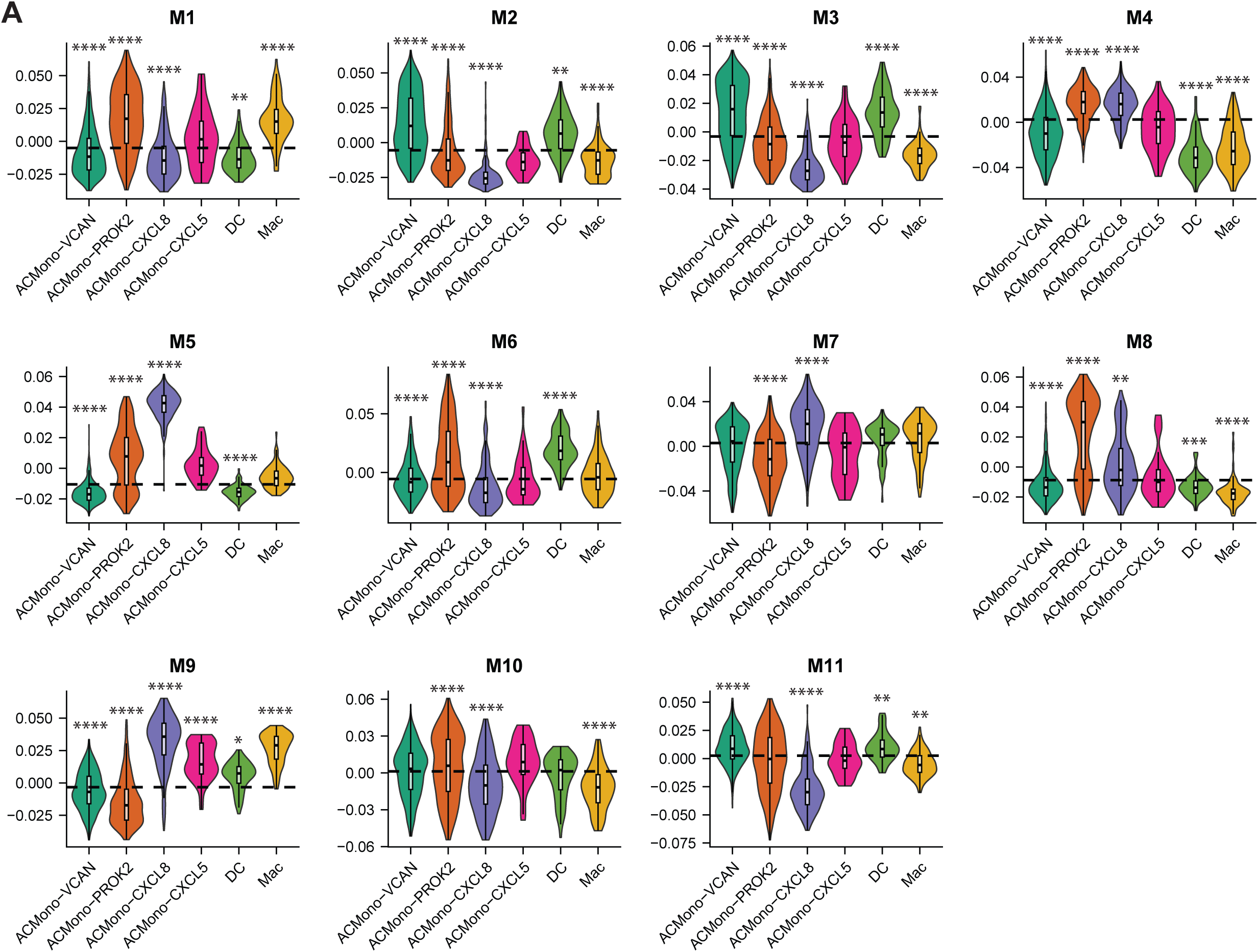
Gene co-expression Modules. 11 monocyte modules differentially expressed across monocyte subtypes.

**Supplementary Figure S8.**
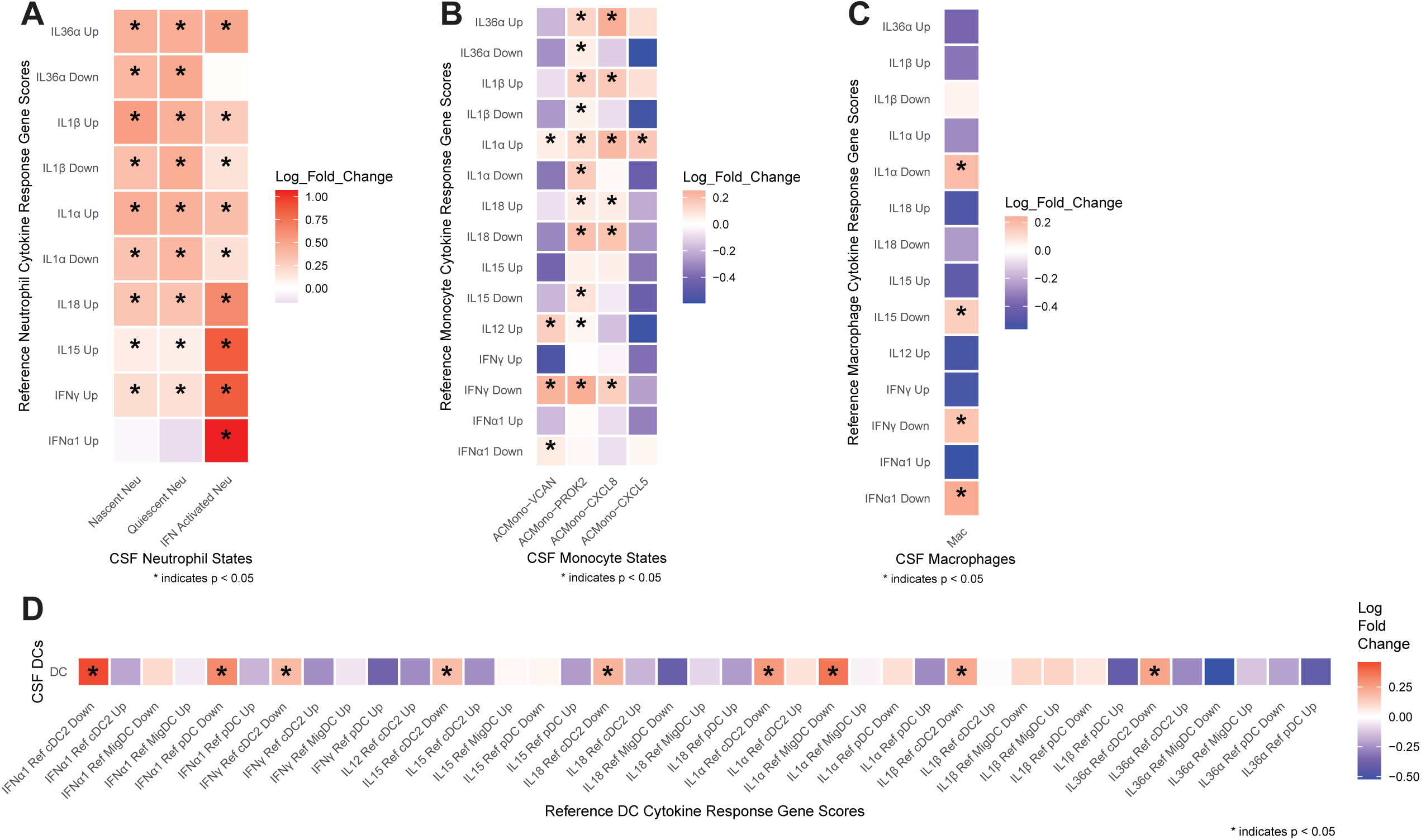
Cytokine Response Scores. Heatmap showing of CSF A. neutrophils, B., monocytes, C. macrophages, and D. dendritic cells showing mean log fold changes of scores based on differential gene expression in response to specific cytokines derived from Immune Dictionary. All interleukin and interferon cytokines eliciting a response in that cell type in Immune Dictionary are plotted.

**Supplementary Table S1.** Marker genes for all cell types.

**Supplementary Table S2.** Marker genes for neutrophil subtypes.

**Supplementary Table S3.** Neutrophil subtype marker genes from prior studies and enrichment scores.

**Supplementary Table S4.** Neutrophil gene co-expression module membership and functional enrichment.

**Supplementary Table S5.** Marker genes for monocyte subtypes.

**Supplementary Table S6.** Monocyte subtype marker genes from prior studies and enrichment scores.

**Supplementary Table S7.** Monocyte gene co-expression module membership and functional enrichment.

**Supplementary Table S8.** Cytokine activity scores using Immune Dictionary signatures and CellChat.

**Supplementary Table S9.** Flow chart of subject recruitment.

## Acknowledgments

P.C. received funding from the Henry M. Jackson Foundation for the Advancement of Military Medicine, Neurocritical Care Society, and Passano Foundation. P.C. and S.A.A. received funding from the Brain Aneurysm Foundation. S.M. was supported by NIH R01 MH052716. The authors thank Flow Cytometry Shared Service of the University of Maryland Marlene and Stewart Greenebaum Comprehensive Cancer Center. This publication was supported by funds through the Maryland Department of Health’s Cigarette Restitution Fund Program – CH-649-CRF and the National Cancer Institute - Cancer Center Support Grant (CCSG) - P30CA134274. Graphical abstract and Figure 1 schematic created in BioRender (https://BioRender.com/q4n1xt9).

---

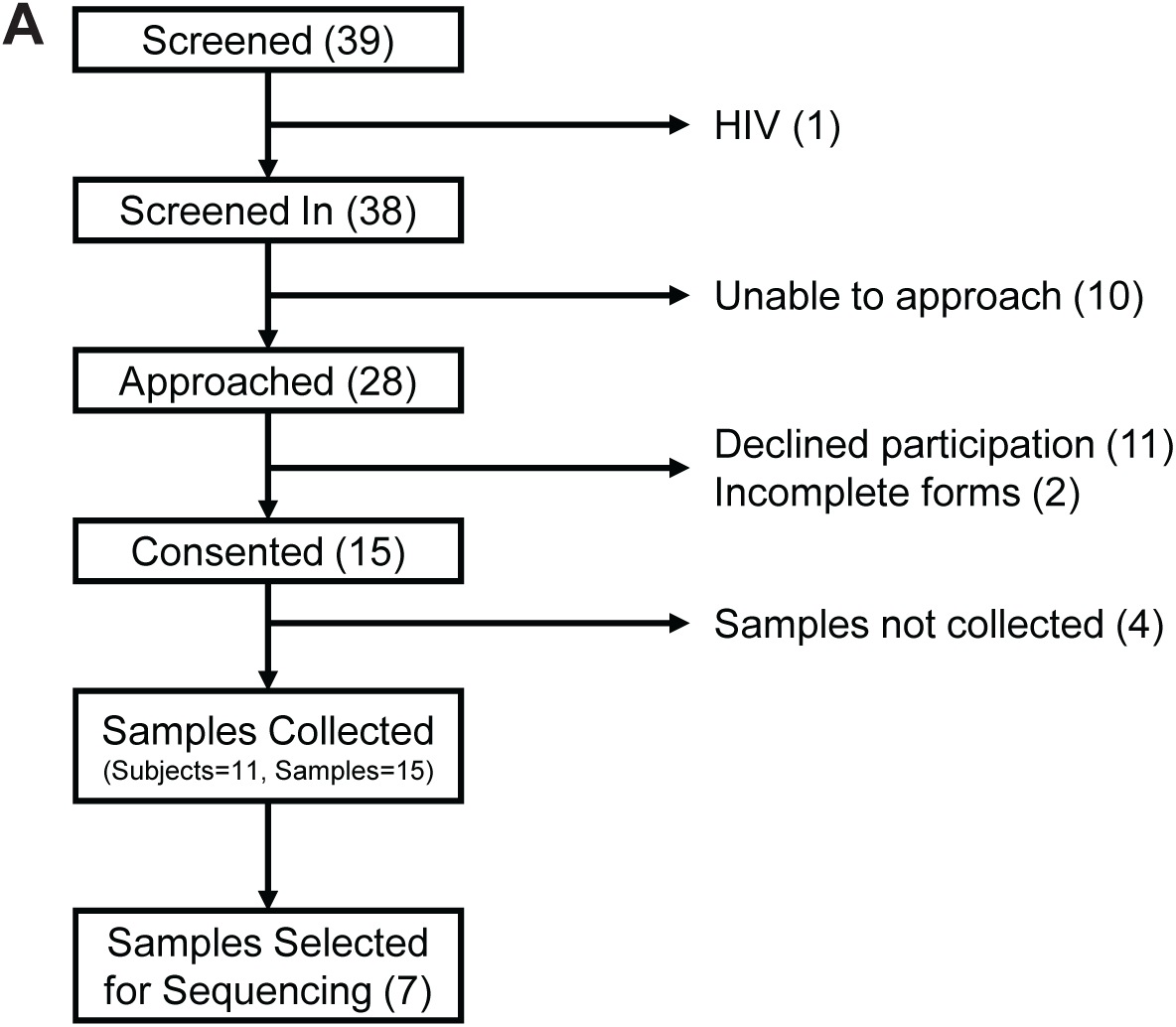

